# A PI3K-WIPI2 positive feedback loop allosterically activates LC3 lipidation in autophagy

**DOI:** 10.1101/2019.12.18.880591

**Authors:** Dorotea Fracchiolla, Chunmei Chang, James H. Hurley, Sascha Martens

**Author notes:** These authors contributed equally (Dorotea Fracchiolla, Chunmei Chang). Corresponding authors: James H. Hurley, Sascha Martens.

## Abstract

Autophagy degrades cytoplasmic cargo by its delivery to lysosomes within double membrane autophagosomes. Synthesis of the phosphoinositide PI(3)P by the autophagic PI 3-kinase complex I (PI3KC3-C1) and conjugation of ATG8/LC3 proteins to phagophore membranes by the ATG12–ATG5-ATG16L1 (E3) complex are two critical steps in autophagosome biogenesis, connected by WIPI2. Here we present a complete reconstitution of these events. On giant unilamellar vesicles (GUVs), LC3 lipidation is strictly dependent on the recruitment of WIPI2, which in turn depends on PI(3)P. Ectopically targeting E3 to membranes in the absence of WIPI2 is insufficient to support LC3 lipidation, demonstrating that WIPI2 allosterically activates the E3 complex. PI3KC3-C1 and WIPI2 mutually promote the recruitment of each other in a positive feedback loop. When both PI 3-kinase and LC3 lipidation reactions were carried out simultaneously, positive feedback between PI3KC3-C1 and WIPI2 led to rapid LC3 lipidation with kinetics similar to those seen in cellular autophagosome formation.

**Summary:** Autophagy requires the synthesis of PI(3)P and the conjugation of LC3 to the phagophore membrane. We reconstituted these two reactions and their coupling by WIPI2, and showed that positive feedback between PI3KC3-C1 and WIPI2 leads to rapid LC3 lipidation by the ATG16L1 complex.

## Introduction

Macroautophagy (hereafter autophagy) is a conserved intracellular degradation process that ensures cellular homeostasis by the degradation of harmful material such as damaged organelles, misfolded proteins and bacterial pathogens. It also ensures survival of cells during starvation (Anding and Baehrecke, 2017; Gomes and Dikic, 2014; Kirkin and Rogov, 2019; Wen and Klionsky, 2016; Zaffagnini and Martens, 2016). These functions are achieved by the sequestration of the cargo material within double membrane vesicles, the autophagosomes, which form *de novo* around the cargo. The precursors of autophagosomes, called phagophores (or isolation membranes) emerge in the cytoplasm as small membrane structures, which capture the cargo as they grow. Upon closure of the phagophores, the structures become autophagosomes, which deliver their cargo for degradation upon fusion with lysosomes (Lamb et al., 2013; Mercer et al., 2018).

Around 40 ATG (AuTophaGy) proteins are required for most forms of autophagy. In particular, these are the ULK1/Atg1 kinase complex, the ATG9 vesicles, the ATG14-containing class III phosphatidylinositol-3 kinase complex I (PI3KC3-C1), the PROPPINs (β-Propellers that bind polyphosphoinositides) and ATG2, as well as the ATG12 and LC3 conjugation machineries (Bento et al., 2016; Hurley and Young, 2017; Mizushima et al., 2011). PI3KC3-C1 phosphorylates phosphatidylinositol at position 3 of the inositol ring and generates phosphatidylinositol-3-phosphate (PI(3)P) on the target membrane, which is essential for autophagy (Itakura and Mizushima, 2009; Matsunaga et al., 2009; Obara et al., 2006). PI(3)P-enriched membranes recruit downstream sensor proteins called PROPPINs or WIPIs (WD-repeat protein interacting with phosphoinositides) (Proikas-Cezanne et al., 2015). WIPIs have a seven bladed β-propeller fold and are a platform for the binding of downstream effector proteins. WIPIs bind PI(3)P and PI(3,5)P2 at two sites through their Phe-Arg-Arg-Gly (FRRG) motifs and increase membrane affinity further using a loop that inserts into the target membrane (Baskaran et al., 2012; Krick et al., 2012; Watanabe et al., 2012). While *S. cerevisiae* has three members of this family, Atg18, Atg21 and Hsv2, humans have 4 orthologues, namely WIPI1-4. Additionally, several isoforms derived from alternative splicing exist for the WIPIs. For example, there are 6 isoforms for WIPI2 (a-e).

WIPIs promote phagophore expansion by the recruitment of the LC3 conjugation machinery (Dooley et al., 2014). In yeast, Atg21 interacts with the coiled-coil domain of Atg16 to recruit the Atg12–Atg5-Atg16 complex to the site of autophagosome formation (Juris et al., 2015). In humans WIPI2b recruits ATG16L1 via a direct interaction with a motif downstream of the coiled-coil domain of ATG16L1 (Dooley et al., 2014). Because the ATG12–ATG5-ATG16L1 complex behaves much like an ubiquitin E3 ligase, we will refer to it from here on as the “E3” complex. The E3 complex subsequently acts to promote the attachment of LC3 proteins to the membrane lipid phosphatidylethanolamine (PE) in a manner analogous to the action of E3-ligases in ubiquitin conjugation reactions (Hanada et al., 2007). The attachment of LC3 to PE, referred to as lipidation, requires the E1-like ATG7 as well as the E2-like ATG3 enzymes, and mediates efficient phagophore elongation and serves to recruit cargo material into autophagosomes (Ichimura et al., 2000; Zaffagnini and Martens, 2016). The E3 complex itself is the product of a ubiquitin-like conjugation machinery wherein the ATG7 and ATG10 proteins conjugate the ubiquitin-like ATG12 to a lysine residue in ATG5 (Mizushima et al., 1998). The ATG12–ATG5 conjugate subsequently non-covalently binds the ATG16L1 protein (Kuma et al., 2002; Mizushima et al., 2003; Mizushima et al., 1999).

A number of the individual steps in autophagy have been studied by *in vitro* reconstitution (Brier et al., 2016), which permits a level of experimental control not possible in cell culture. This minimalist approach can reveal the inherent activities of individual components stripped of the complexity of the cellular context. More sophisticated reconstitutions including multiple purified components and reactions can restore some of the physiological context and complexity while maintaining maximal experimental control. As part of a larger effort to reconstitute mammalian autophagosome biogenesis from start to finish using giant unilamellar vesicles (GUVs) as a model, we simultaneously reconstituted the PI3KC3-C1 lipid phosphorylation and E3 promoted LC3 lipidation reactions as coupled by WIPI2. These experiments uncovered the existence of a positive feedback loop involving the PI3KC3-C1 and WIPI2, wherein the two factors mutually enhance their membrane recruitment. In addition, we found that WIPI2 does not merely recruit the E3 to the membrane, but allosterically activates it as well. Working in combination, these two effects propel the rapid LC3 lipidation on synthetic GUV membranes on a timescale similar to that seen in cells.

## Results

### Reconstitution of PI(3)P- and WIPI-dependent LC3 lipidation

We expressed and purified the recombinant human E3 complex by co-expressing its three subunits ATG12, ATG5 and ATG16L1 (isoform ß) together with ATG7 and ATG10 in insect cells (**Figure S1A**). We assessed the oligomeric state of the purified complex in solution by static light scattering (SLS) and detected a monodisperse population of dimeric complexes, where the experimentally determined molecular weight corresponds to 2.90 x 10^5^ Da ± 0.8 %, essentially equal to the computed mass of two 1.48 x10^5^ Da monomers (**Figure S1B**). Thus, the purified E3 complex exists as a dimer composed of two copies of each subunit ATG5, ATG12, ATG16L1 (**Figure S1C**). We assessed the ability of the recombinant E3 complex to catalyze LC3 lipidation. We co-incubated PE-containing SUVs with ATG7, ATG3, and LC3BΔ5C, with the latter lacking the 5 C-terminal residues in order to expose the Gly required for lipidation (LC3B-I) (see all purified components of the machinery in **Figure S1D**). Consistent with the observation by (Lystad et al., 2019), we found that the presence of the E3 complex promoted LC3B lipidation (LC3B-II) (**Figure 1A**). We also found that the presence of ATG16L1 was required for the promotion of LC3B-II formation (**Figure 1A**). Lipidation was far more efficient with dioleoyl (DO) lipids as compared to palmitoyl-oleoyl (PO) lipids (**Figure S1E, F**). LC3-II conversion was more than 50% complete with DO lipids after 30 min, while a comparable degree of conversion required overnight incubation with PO lipids. This effect cannot be explained by preferential membrane binding by the E3 complex (Dudley et al., 2019; Lystad et al., 2019), as PO and DO lipids are bound equally (**Figure S1G, H**). These data, which are consistent with a high proportion of unsaturated lipids in autophagosomal membranes (Schütter et al., 2020), suggest that the presence of two rather than one unsaturated tail in the lipids of the membrane substrates increases the flexibility of the membrane, facilitating structural rearrangements necessary for LC3 lipidation.

**Figure 1.**
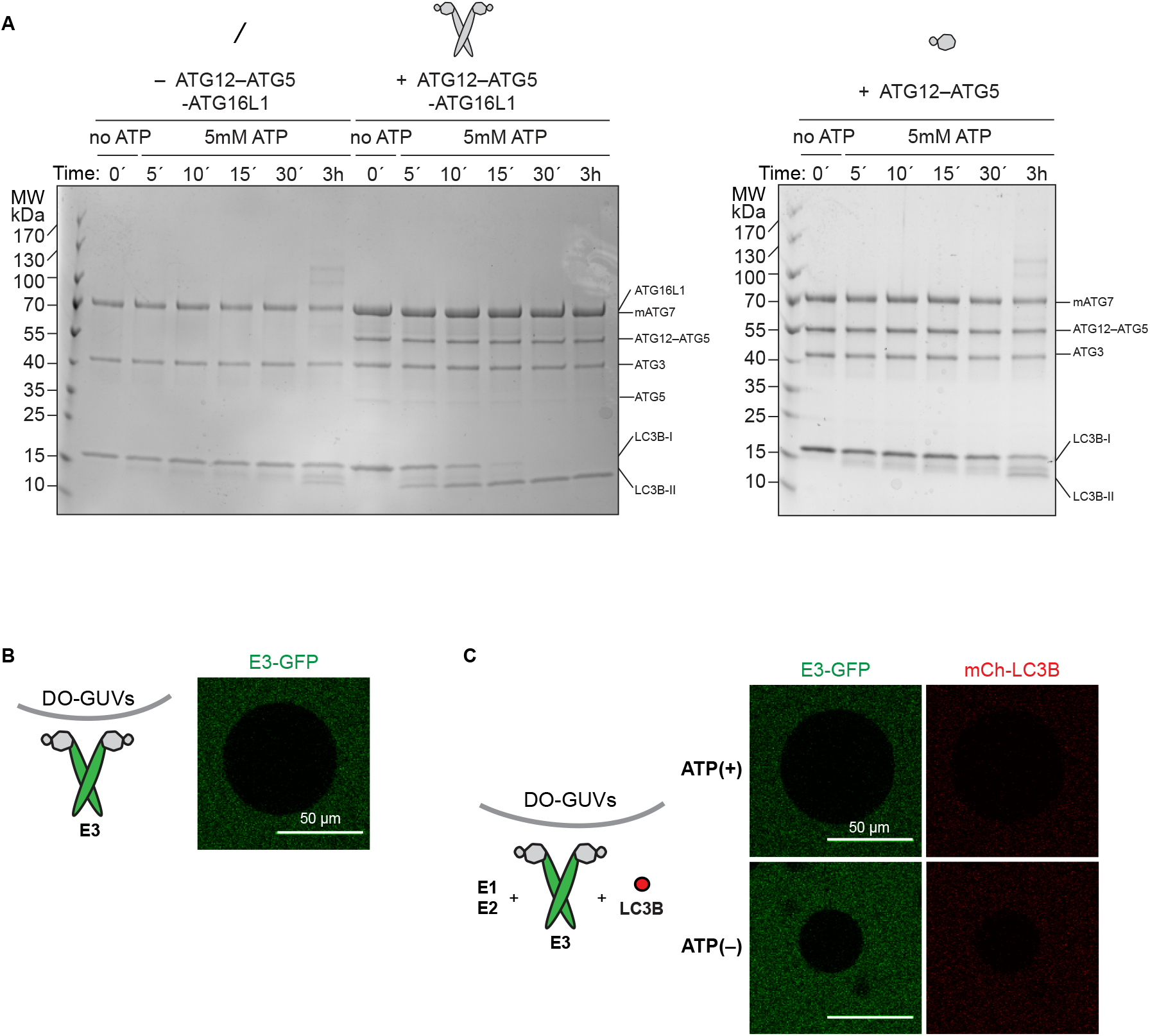
The LC3 lipidation machinery is active on SUVs but not on GUVs. **A** *In vitro* LC3B lipidation assay on DO-SUVs. ATG7, ATG3 and LC3B were mixed with SUVs (65% PC: 15% liver PI: 20% PE), in the absence (left) or presence of the ATG12-ATG5-ATG16L1 (E3) complex (1 μM) (middle) or the ATG12-ATG5 conjugate (1 μM) (right) and incubated at 37°C in the presence of MgCl_2_/ATP. Samples taken at the indicated time points were loaded on a 4-15% SDS PAGE. **B** The E3-GFP complex (0.5 μM) was added to DO-GUVs (65% PC: 15% liver PI: 20% PE). Scale bars, 50 μm. **C** The E3-GFP (0.5 μM), ATG7, ATG3 and mCherry-LC3B were added to GUVs (65% PC: 15% liver PI: 20% PE), in the presence of MgCl_2_/ATP. Representative confocal micrographs are shown. Scale bars, 50 μm.

In cells, efficient membrane recruitment of the E3 complex during autophagosome formation depends on the presence of WIPI2 (Dooley et al., 2014). In order to recapitulate this step, we employed GUVs instead of SUVs because they are less curved, rendering membrane binding more stringent. In the absence of WIPI2d, we observed no E3 recruitment to GUVs bearing the same lipid composition as the DO-SUVs used in the previous assays (compare **Figure S1H** and **Figure 1B**). The addition of ATG7, ATG3, mCherry-LC3BΔ5C and ATP did not result in any detectable mCherry-LC3B lipidation on GUVs (**Figure 1C**).

To test the ability of WIPIs to activate LC3 lipidation, we expressed and purified the three human WIPI proteins WIPI2d, WIPI3 and WIPI4 isoform 1. We assessed their ability to interact with the E3 complex in a microscopy-based bead interaction assay (**Figure 2A, S2A**). mCherry-WIPI2d was specifically recruited to beads coated with E3-GFP (**Figure 2A**). We also observed that the E3-GFP was robustly recruited to beads coated with GST-mCherry-WIPI3, but not GST-mCherry-WIPI4 (**Figure S2A**). Thus, the E3 complex directly binds to WIPI2d and WIPI3. We then tested the ability of WIPI2d to recruit the E3 to GUVs. We added the E3-GFP and mCherry-WIPI2d to PI(3)P-containing GUVs and compared the GFP signal intensity to conditions lacking mCherry-WIPI2d (**Figure 2B**) or PI(3)P (**Figure S2B**). The E3 was robustly and specifically recruited to PI(3)P-containing GUVs in a WIPI2d dependent manner (**Figure 2B**). When 15% PI was used instead of 5% PI(3)P, no WIPI2d and E3 recruitment was observed, despite the equivalent negative charge of the membrane (**Figure S2B**).

**Figure 2.**
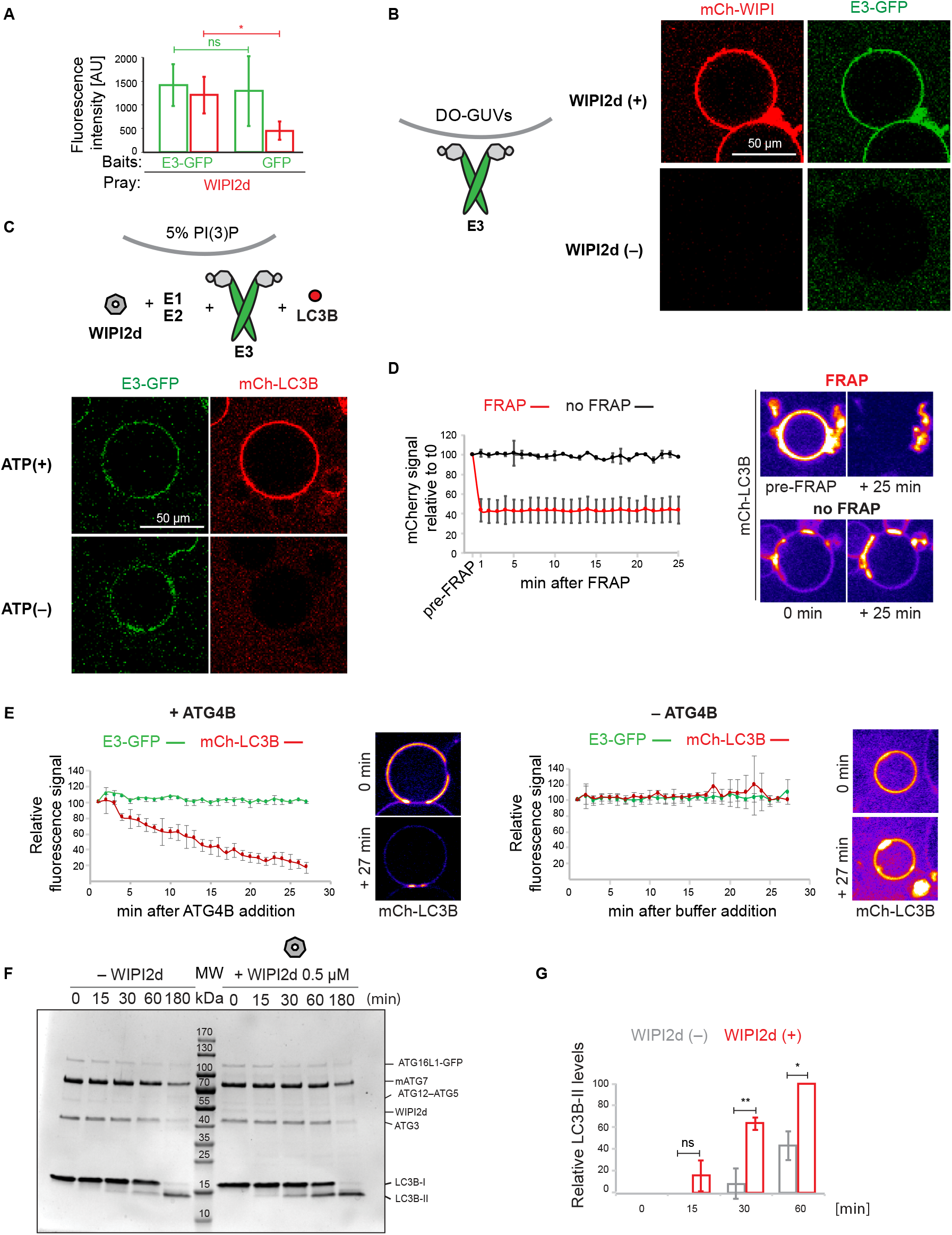
WIPI2d both recruits and allosterically activates E3 for LC3 lipidation. **A** Quantification of the mCherry-WIPI2d signal intensity (red bars) measured on GFP-Trap beads coated with either GFP or GFP-tagged E3 (means ± SDs are shown; N=94 (E3-GFP) or 80 (GFP)). **B** The E3-GFP (0.5 μM) was added to DO-GUVs containing 75% PC: 5% PI(3)P: 20% PE in the absence or presence of mCherry-WIPI2d (0.5μM). Scale bars, 50 μm. **C** The E3-GFP (0.1 μM) was co-incubated with WIPI2d (0.5 μM), mCherry-LC3B and the lipidation machinery on DO-GUVs (75% PC: 5% PI(3)P: 20% PE) in the presence of MgCl_2_/ATP. Scale bars, 50 μm. **D** FRAP experiment on GUVs after lipidation in the presence of ATP as conducted in **C**. A quantification is shown (means ± SDs are shown; N FRAP=3, N no FRAP=2) together with representative images of the two conditions, at time 0 and 25 minutes after the photobleaching. **E** De-lipidation reactions on GUVs treated as in **C**, in the presence of ATP. CIP/ATG4B (left) or buffer (right) were added to the wells and imaging was conducted for the indicated time. Quantification of the mCherry-LC3B and E3-GFP signals overtime is shown (means ± SDs are shown; CIP/ATG4B =31, N buffer=6) together with representative images of the two conditions at time 0 and 27 minutes after the addition. **F** *In vitro* LC3B lipidation assay on DO-SUVs in the absence or presence of WIPI2d. ATG7, ATG3, E3-GFP (0.1 μM) and LC3B are mixed with DO-SUVs (75% PC: 5% PI(3)P: 20% PE), either in the absence (left) or in the presence of 0.5 μM WIPI2d (right) and incubated at 37°C with MgCl_2_/ATP. Samples were taken at the indicated timepoints and loaded on a 415% SDS PAGE. **G** Quantification of three independent experiments is shown as relative LC3B-II levels at each time point (means ± SDs are shown; N = 3). P-values were calculated using Student’s *t*-test (p≥0.5: (ns); 0.01<p<0.05 (*); 0.001<p<0.01: (**)).

We assessed whether the recruitment of the E3 to PI(3)P-containing GUVs by WIPI2d elicited LC3B lipidation. mCherry-LC3B was efficiently lipidated to these GUVs in an ATP, WIPI2d and PI(3)P-dependent manner (**Figure 2C, S2C**). In order to further confirm that the mCherry signal observed on these GUVs was due to covalent attachment of LC3B to membranes, we performed Fluorescence Recovery After Photobleaching (FRAP) experiments (**Figure 2D**) (Fracchiolla et al., 2016). The mCherry-LC3B signal did not recover over the time period imaged suggesting that mcherry-LC3B is covalently linked to the membrane. Moreover, the addition of the ATG4B protease, which removed LC3B from the membrane resulted in loss of membrane associated fluorescence further corroborating that LC3B is lipidated on the GUV membrane (**Figure 2E**). Thus, the GUV system recapitulates the strict dependence of E3 recruitment and LC3B lipidation on PI(3)P and WIPI2 observed in cells (Dooley et al., 2014).

### WIPI2d both recruits and allosterically activates E3 for LC3 lipidation

Given the dramatic stimulatory effect of WIPI2d on LC3B lipidation in the GUV system (**Figure 2C**), we asked whether WIPI2d could activate the E3 complex beyond merely recruiting it to the membrane. We observed that the lipidation occurred more efficiently in the presence of WIPI2d (**Figure 2F, G**). Importantly, in the SUV system WIPI2d did not significantly increase membrane binding by the E3 as assessed by sedimentation assays (**Figure S3A)** and bead-based recruitment assays (**Figure S3B, C**). We therefore conclude that WIPI2d must have an effect on the LC3B lipidation machinery that goes beyond simply recruiting E3 to the membrane.

In yeast, the WIPI ortholog Atg21 recruits Atg16 to the phagophore assembly site (PAS)(Juris et al., 2015). Atg21 binds to a DE motif (D101, E102) within the coiled-coil of Atg16, while WIPI2b binds to an EE-containing motif C-terminal to the extended coiled-coil domain (E226, E230)(Dooley et al., 2014) (**Figure 3A, B**). Fortuitously, the intra-coiled coil motif of yeast Atg16 is also present in human ATG16L1, and corresponds to residues D164,E165. We found that yeast Atg21 can directly bind the human E3 complex (**Figure 3D**). This led us to ask if E3 activation was unique to human WIPI2 or common to any ATG16- and PI(3)P-binding WIPI ortholog. Atg21 was added to LC3 lipidation reactions using SUVs. Atg21 did not significantly enhance LC3 lipidation (**Figure 3E, F**). Thus, the stimulatory effect of WIPI2d as opposed to Atg21 cannot be attributed to enhanced membrane binding of the E3 in the presence of WIPI2d because the membrane binding by the E3 was not significantly different under the conditions tested (**Figure S3A, B, C**).

**Figure 3.**
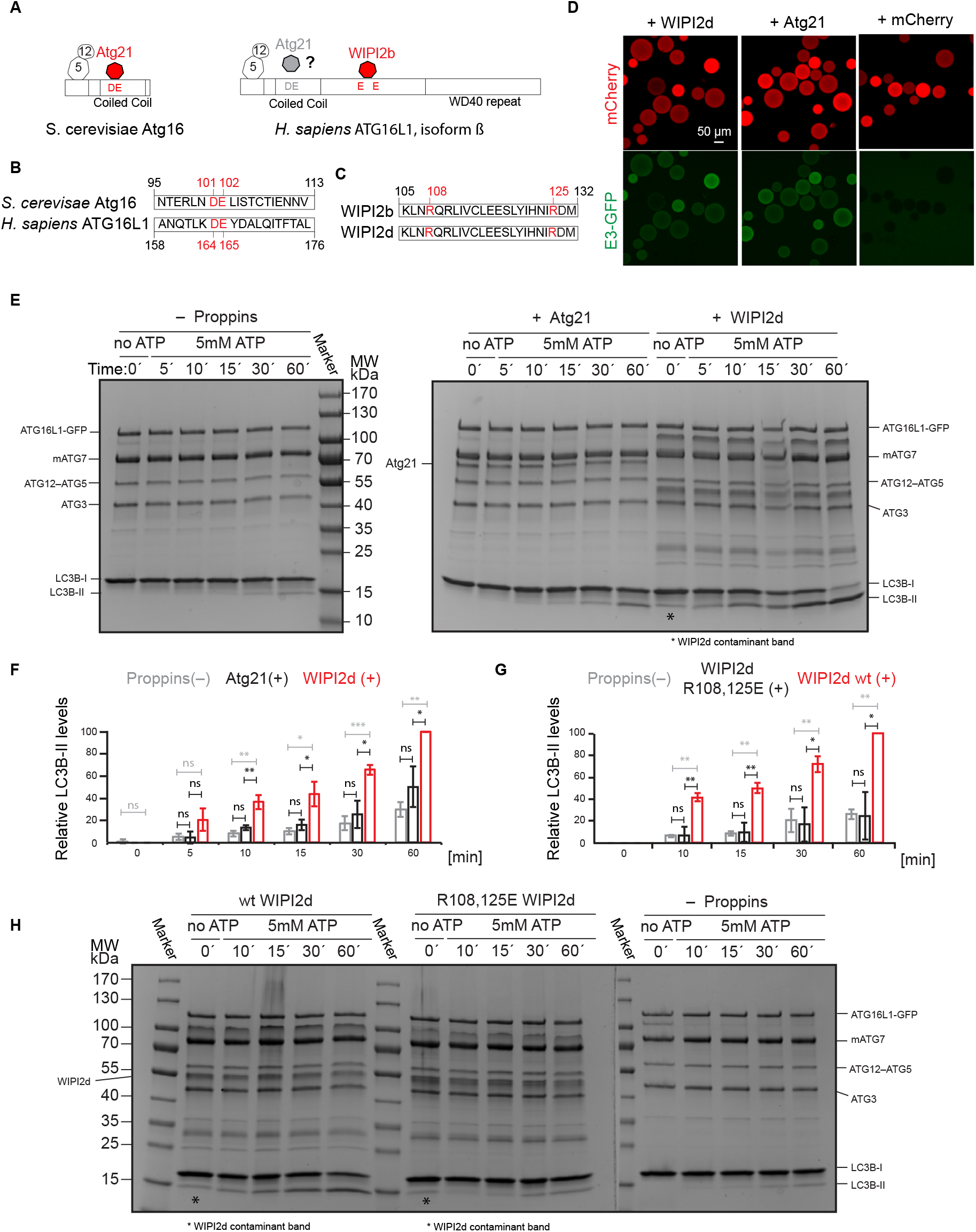
WIPI2d specifically promotes LC3 lipidation. **A** Scheme showing the domain organization of *S. cerevisiae* Atg16 and *H. sapiens* ATG16L1 (isoform β) and their interactors. **B** Alignment of the Atg16 and *H. sapiens* ATG16L1 (isoform β) protein sequences spanning a region around the D101, E102 of Atg16 (Atg21 binding site) and residues D164, E165 in ATG16L1. **C** Alignment of the *H. sapiens* WIPI2 isoforms b and d spanning the region surrounding the ATG16L1 binding site including R108 and R125. **D** Microscopy-based bead protein interaction assay with RFP-Trap beads coated with mCherry-WIPI2d, mCherry-Atg21 or mCherry as baits and incubated with 5 μM E3-GFP as pray. Representative confocal micrographs are shown. Scale bars, 50 μm. **E, F** DO-SUVs (75% PC: 5% PI(3)P: 20% PE) were incubated with ATG7, ATG3, E3-GFP (0.5 μM) and LC3B in the presence of no PROPPIN, WIPI2d or Atg21 (both 2.5μM). Samples taken at the indicated time points were loaded on a 4-15% SDS PAGE. The quantification is shown in **Figure 3F** as relative LC3B-II levels for each timepoint (means ± SDs are shown; N = 3). **G, H** DO-SUVs (75% PC: 5% PI(3)P: 20% PE) were incubated with ATG7, ATG3, E3-GFP (0.5 μM) and LC3B in the presence of wt WIPI2d, R108,125E mutant WIPI2d (at 2.5μM) or no PROPPINs. Samples taken at the indicated time points were loaded on a 4-15% SDS PAGE. The quantification is shown in **Figure 3G** as relative LC3B-II levels for each timepoint (means ± SDs are shown; N = 3). P-values were calculated using Student’s *t*-test (p≥0.5 (ns); 0.01<p<0.05: (*); 0.001<p<0.01: (**); p<0.001 (***)).

We took advantage of the previously characterized ATG16L1 binding site in WIPI2b identified in (Dooley et al., 2014). Sequence alignment of the WIPI2b and d isoforms showed that the positively charged residues R108 and R125 comprising the binding site are present in both isoforms (**Figure 3C**). The R108,125E mutant was defective in binding to the E3 complex on GUVs (**Figure S3D**) and lost the promoting effect on LC3B lipidation (**Figure 3G, H, S3E**), though still retaining its ability to bind PI(3)P membranes (**Figure S3F**). We conclude that WIPI2d has a potent activating effect on LC3B lipidation depending on its ability to recruit the E3 to the membrane via the ATG16L1 subunit. Beyond this, it further activates LC3B lipidation likely due to an allosteric effect on the E3.

### PI3KC3-C1 supports LC3B lipidation on unsaturated flat membranes

During autophagosome nucleation, the PI3KC3-C1 complex translocates to the endoplasmic reticulum (ER) to generate PI(3)P to recruit downstream factors (Axe et al., 2008; Matsunaga et al., 2010). It was previously reported that PI3KC3-C1 is active only on high curvature membranes and has no measurable activity on GUV membranes (Rostislavleva et al., 2015). The ER membrane contains both flat and highly curved membranes, and much of the omegasome domain of the ER involved in autophagy is flat on a molecular scale. The membrane of the ER is less densely packed than post-Golgi compartments (Bigay and Antonny, 2012; Vanni et al., 2014), and we reasoned that highly unsaturated lipid mixtures, also found in autophagosomal membranes (Schütter et al., 2020), might therefore support PI3KC3-C1 activity on the flat membrane of GUVs.

We tested membrane binding and the enzymatic activity of PI3KC3-C1 on GUVs composed of lipids with a headgroup composition resembling the ER (65% PC: 20% PE: 5% PS: 10% PI) but with different hydrophobic tails. PI3KC3-C1 failed to bind to GUV membranes made with the brain lipid extract previously used (Rostislavleva et al., 2015) (**Figure S4A, B**). Using a mCherry-FYVE (Fab1, YOTB, Vac1, and EEA1) domain as a probe for PI(3)P, minimal activity of PI3KC3-C1 was observed on these GUVs (**Figure S4A, C**). In contrast, PI3KC3-C1 strongly bound to and robustly produced PI(3)P on GUVs composed of lipids with two unsaturated tails (DO), but not of only one unsaturated (PO) tail (**Figure 4A, B, S4**). Little FYVE domain recruitment was seen on GUVs composed of DO lipids in the absence of PI3KC3-C1 (**Figure S4A, C**). These data show that PI3KC3-C1 is active on flat membranes with unsaturated lipids, suggesting that ER-like membranes with loose lipid packing are good substrates for PI3KC3-C1 even when they are flat.

**Figure 4.**
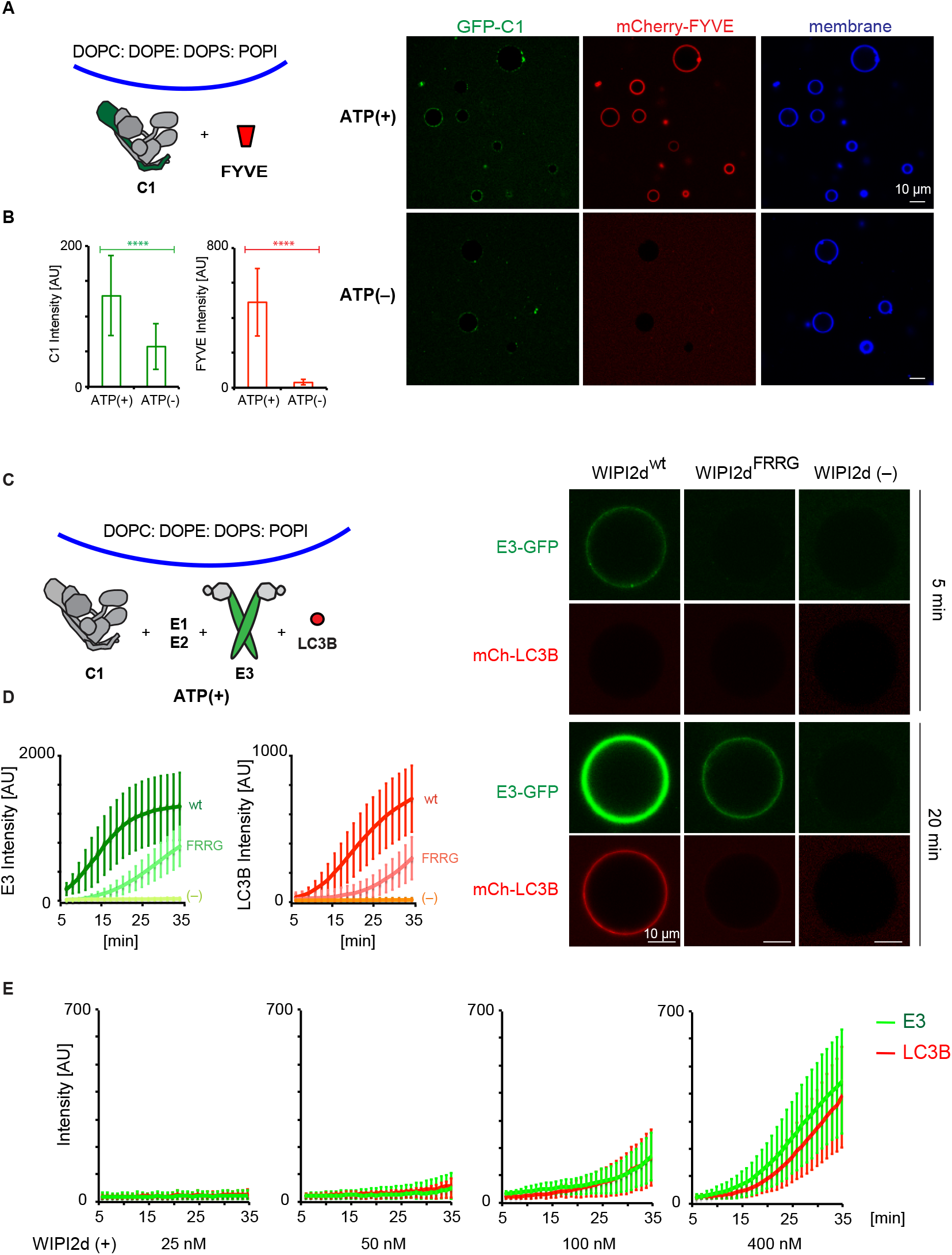
PI3KC3-C1supports LC3B lipidation on unsaturated flat membranes. The schematic drawing illustrates the reaction setting. Colors indicate fluorescent protein fused components. Components in gray are not labeled but are present in the reaction mix. **A** Representative confocal images of GUVs showing the membrane binding of the PI3KC3-C1 and FYVE domain. GFP-tagged PI3KC3-C1 (200 nM) and mCherry-tagged FYVE (1 μM) were incubated with DO-GUVs (64.8% PC: 20% PE: 5% PS: 10% POPI: 0.2% Atto647DOPE) in the presence or absence of ATP/Mn^2+^ (50 μM/1 mM) at room temperature. Images were taken after 30 min incubation. Scale bars, 10 μm. **B** Quantification of the relative intensities of PI3KC3-C1 (green bars) and FYVE domain (red bars) on GUV membranes in **A** (means ± SDs; N = 50). P-values were calculated using Student’s *f*-test (p≥0.5 (ns); 0.01<p<0.05 (*)). **C** Representative confocal images of GUVs showing E3 binding and LC3B lipidation. mCherry-tagged LC3B was incubated with GUVs in the presence of PI3KC3-C1 (0.1 μM), WIPI2d (0.4 μM or none) or WIPI2d FRRG mutant (0.4 μM), E3-GFP, ATG7, ATG3, and ATP/Mn^2+^ (50 μM/1 mM). Images taken at indicated time points were shown. Scale bars, 10 μm. **D** Quantitation of the kinetics of E3 recruitment and LC3B lipidation on the membrane from individual GUV tracing in **C** (means ± SDs; N = 53 (wt), 45 (FRRG), 52 (-)). **E** Quantitation of the kinetics of E3 recruitment and LC3B lipidation on the membrane from individual GUV tracing (means ± SDs; N = 26 (25 nM); 40 (50 nM); 37 (100 nM); 32 (400 nM)). mCherry-LC3B was incubated with PI3KC3-C1 (0.1 μM), E3-GFP, ATG7 and ATG3 in the presence of WIPI2d with difference concentration.

We tested whether PI3KC3-C1 could support LC3B lipidation on GUV membranes with DO lipids *via* the WIPI2d – E3 axis. Upon addition of WIPI2d, the E3 and the LC3 conjugation machinery (ATG7, ATG3, mCherry-LC3BΔ5C), PI3KC3-C1 robustly triggered membrane recruitment of the E3-GFP complex and subsequent mCherry-LC3B lipidation (**Figure 4C, D**) Consistent with expectation, the E3 was not detectably recruited to the GUV membrane in the absence of WIPI2d (**Figure 4C, D**). Mutation of the conserved PI(3)P binding FRRG motif significantly reduced E3 membrane recruitment and LC3 lipidation (**Figure 4C, D**). The effect of WIPI2d on the PI3KC3-C1–induced E3 membrane association and LC3B lipidation was dose-dependent (**Figure 4E**). We analyzed the effect of WIPI3 on LC3B lipidation in the *in vitro* system and found that WIPI3 also mediated LC3B lipidation on GUV membranes (**Figure S5**), suggesting a potential role of WIPI3 in autophagosome formation in cells. In combination, the data obtained using the reconstituted system showed that PI3KC3-C1 stimulates LC3 lipidation on GUV membranes in a WIPI-dependent manner.

### Positive feedback between PI3KC3-C1 and WIPI2d promotes LC3B lipidation

Since PI3KC3-C1 strongly promotes WIPI2-dependent LC3B lipidation (**Figure 4C, D**), we asked if it was solely attributable to its PI 3-kinase enzyme activity or if the PI3KC3-C1 also had additional roles. To elucidate the role of PI3KC3-C1 more precisely, we assayed how much PI(3)P was produced by PI3KC3-C1 on GUVs by comparing FYVE domain recruitment to GUVs with 10% (mol fraction) PI(3)P in the absence of C1 or GUVs with 10% PI in the presence of C1, respectively. We found that the PI3KC3-C1 produced about 2% PI(3)P in 30 min (**Figure 5A, B**). We then measured WIPI2d recruitment to these two GUVs and surprisingly found that WIPI2d was recruited much faster than the FYVE domain to the 10% PI GUVs in the presence of PI3KC3-C1 despite showing lower affinity than the FYVE domain to 10% PI(3)P GUVs (**Figure 5A, B**). The WIPI2d FRRG mutant was not detectably recruited to the either of these two GUVs (**Figure 5A, B**).

**Figure 5.**
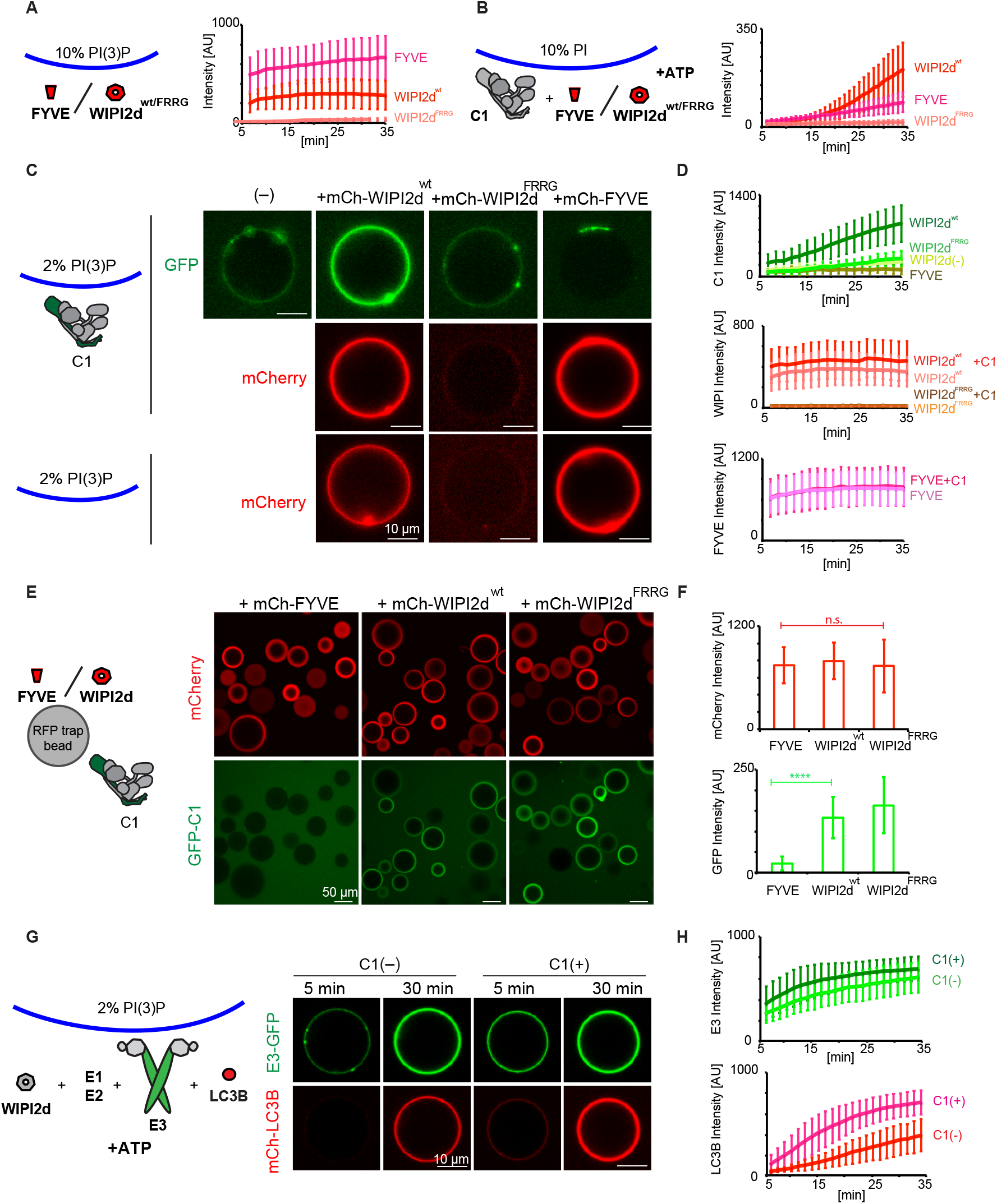
Positive feedback between PI3KC3-C1 and WIPI2d promotes LC3B lipidation. **A** Quantitation of the kinetics of FYVE domain or WIPI2d recruitment to the membrane from individual GUV tracing (means ± SDs; N = 45 (FYVE); 54 (WIPI2d^wt^); 40 (WIPI2d^FRRG^)). mCherry-FYVE, mCherry-WIPI2d, or mCherry-WIPI2d FRRG mutant (0.5 μM) wasincubated with 10% PI(3)P DO-GUVs (64.8% PC: 20% PE: 5% PS: 10% PI(3)P: 0.2% Atto647 DOPE). **B** Quantitation of the kinetics of FYVE domain or WIPI2d recruitment to the membrane from individual GUV tracing (means ± SDs; N = 53 (FYVE); 64 (WIPI2d^wt^); 66 (WIPI2d^FRRG^)). mCherry-FYVE, mCherry-WIPI2d, or mCherry-WIPI2d FRRG mutant (0.5 μM) and PI3KC3-C1 (0.1 μM) were incubated with 10% PI DO-GUVs (64.8% PC: 20% PE: 5% PS: 10% POPI: 0.2% Atto647 DOPE) in the presence of ATP/Mn^2+^ (50 μM/1 mM). **C** Representative confocal images showing the membrane binding of the PI3KC3-C1 complex, WIPI2d, WIPI2d FRRG mutant or FVYE domain. GFP-tagged PI3KC3-C1 (0.1 μM) was incubated with 2% PI(3)P DO-GUVs (72.8% PC: 20% PE: 5% DOPS: 2% PI(3)P: 0.2% Atto647 DOPE) in the absence or presence of 250 nM mCherry tagged-WIPI2d, WIPI2d FRRG mutant, or FYVE domain, respectively (upper two panels). 250 nM WIPI2d, WIPI2d FRRG mutant, or FYVE domain was incubated with 2% PI(3)P GUVs in the absence of PI3KC3-C1 (bottom panel). Scale bars, 10 μm. **D** Quantitation of the kinetics of PI3KC3-C1, WIPI2d, WIPI2d FRRG mutant, or FYVE domain recruitment to the membrane from individual GUV tracing in **C** (means ± SDs; N = 47 (C1 alone); 48 (C1+WIPI2^wt^); 40 (C1+WIPI2^FRRG^); 33 (C1+FYVE); 42 (WIPI2^wt^ alone); 43 (WIPI2^FRRG^ alone); 42 (FYVE alone)). **E** Microscopy-based bead protein interaction assay with RFP-Trap beads coated with mCherry-FYVE, mCherry-WIPI2d, or mCherry-WIPI2d FRRG mutant as baits and incubated with 0.1 μM GFP-PI3KC3-C1 as pray. Representative confocal micrographs are shown. Scale bars, 50 μm. **F** Quantification of the GFP-PI3KC3-C1 signal intensity (green bars) measured on RFP-Trap beads coated with mCherry-FYVE, mCherry-WIPI2d or mCherry-WIPI2d FRRG mutant (means ± SDs are shown; N = 75). P-values were calculated using Student’s *t*-test (p≥0.5: (ns); 0.01<p<0.05: (*)). **G** Representative confocal images of GUVs showing E3 recruitment and LC3B lipidation. mCherry-LC3B was incubated with 2% PI(3)P GUVs in the presence of PI3KC3-C1 (0.1 μM or none), WIPI2d (0.25 μM), E3-GFP, ATG7, ATG3, and ATP/Mn^2+^. Scale bars, 10 μm. **H** Quantitation of the kinetics of E3 recruitment and LC3B lipidation on the membrane from individual GUV tracing in G (means ± SDs; N = 20 (C1-); 31 (C1+)).

To determine if the enzymatic activity of the PI3KC3-C1 was required for the enhanced recruitment of WIPI2d compared to the FYVE domain, we generated GUVs containing 2% PI(3)P but no PI. We first analyzed the binding of PI3KC3-C1 itself to these GUV membranes, and only trace binding of PI3KC3-C1 to membrane was observed (**Figure 5C, D**). However, in the presence of WIPI2d, but not the WIPI2d FRRG mutant or FYVE domain, an increased amount of PI3KC3-C1 was recruited to the membrane (**Figure 5C, D**). We observed that more WIPI2d was recruited to these GUVs in the presence of PI3KC3-C1 (**Figure 5C, D**), which suggests that PI3KC3-C1 and WIPI2d cooperatively bind to membranes. We further observed that GFP tagged PI3KC3-C1 was specifically recruited to beads coated with WIPI2d, but not the FYVE domain (**Figure 5E, F**). These data imply that the two might form a stable physical complex on membranes, which requires PI(3)P to anchor WIPI2d, but is otherwise separate from the enzymatic activity PI3KC3-C1.

We went on to test LC3B lipidation in the absence or presence of PI3KC3-C1 on GUVs containing PI(3)P but not PI. Consistent with the observation that PI3KC3-C1 and WIPI2d mutually enhance their membrane binding, the addition of PI3KC3-C1 to PI(3)P-containing GUVs increased the efficiency of E3 recruitment and subsequent LC3B lipidation, even though none of the PI3KC3-C1 substrate, PI, was present (**Figure 5G, H**). These data indicate that the activities of the PI3KC3-C1 in autophagy go beyond simply generating PI(3)P. PI3KC3-C1 also cooperatively enhances membrane binding by WIPI2d. By recruiting more PI3KC3-C1, WIPI2d increases PI(3)P production and creates a positive feedback loop leading to even more of its own recruitment. The rapid increase in the amount of WIPI2d on the membrane in turn provides a means to rapidly recruit E3 and allosterically promote LC3 lipidation.

## Discussion

Here we present a full reconstitution of the events during autophagosome formation from PI(3)P production by the PI3KC3-C1 to LC3 lipidation, involving the activity of 13 polypeptides (VPS34, VPS15, BECN1, ATG14, WIPI2d, ATG16L1, ATG12, ATG10, ATG7, ATG5, ATG3, ATG4B and LC3B) in the context of SUV and GUV membranes. The GUV system recapitulated the strict requirement of LC3B lipidation on the presence of PI3KC3-C1 and WIPI2d as demonstrated in cells (Axe et al., 2008; Dooley et al., 2014; Itakura et al., 2008; Itakura and Mizushima, 2010; Matsunaga et al., 2009; Zhong et al., 2009). We showed that a completely defined system of purified proteins and lipids mimics a complex and centrally important pair of coupled steps in autophagosome biogenesis. Moreover, we uncovered mutually stimulatory effects that would have been difficult, if not impossible, to resolve with cell-based assays alone.

We discovered the existence of a positive feedback loop between the PI3KC3-C1 and WIPI2d, which mutually enhances their recruitment to the membrane (**Figure 6**). Our data suggest that once PI3KC3-C1 is localized to the site of autophagosome formation and produces PI(3)P, the PI(3)P recruits the WIPIs, which in turn recruit further PI3KC3-C1 complexes, resulting in rapid PI(3)P production and WIPI recruitment. This translates into an efficient recruitment and activation of the LC3 lipidation machinery. Mechanistically, the existence of this feedback loop is supported by our evidence that PI3KC3-C1 complex and WIPI2d physically interact with one another on membranes, and most probably do so during autophagy induction in cells. This implies that PI3KC3-C1 functions, at least at the earliest onset of autophagosome biogenesis, stoichiometrically with respect to WIPI2. Both PI3KC3-C1 and WIPI2d can additionally insert parts of themselves into the membrane (Baskaran et al., 2012; Fan et al., 2011; Rostislavleva et al., 2015; Chang et al., 2019), thus these two factors could modulate local membrane properties to enhance binding.

**Figure 6.**
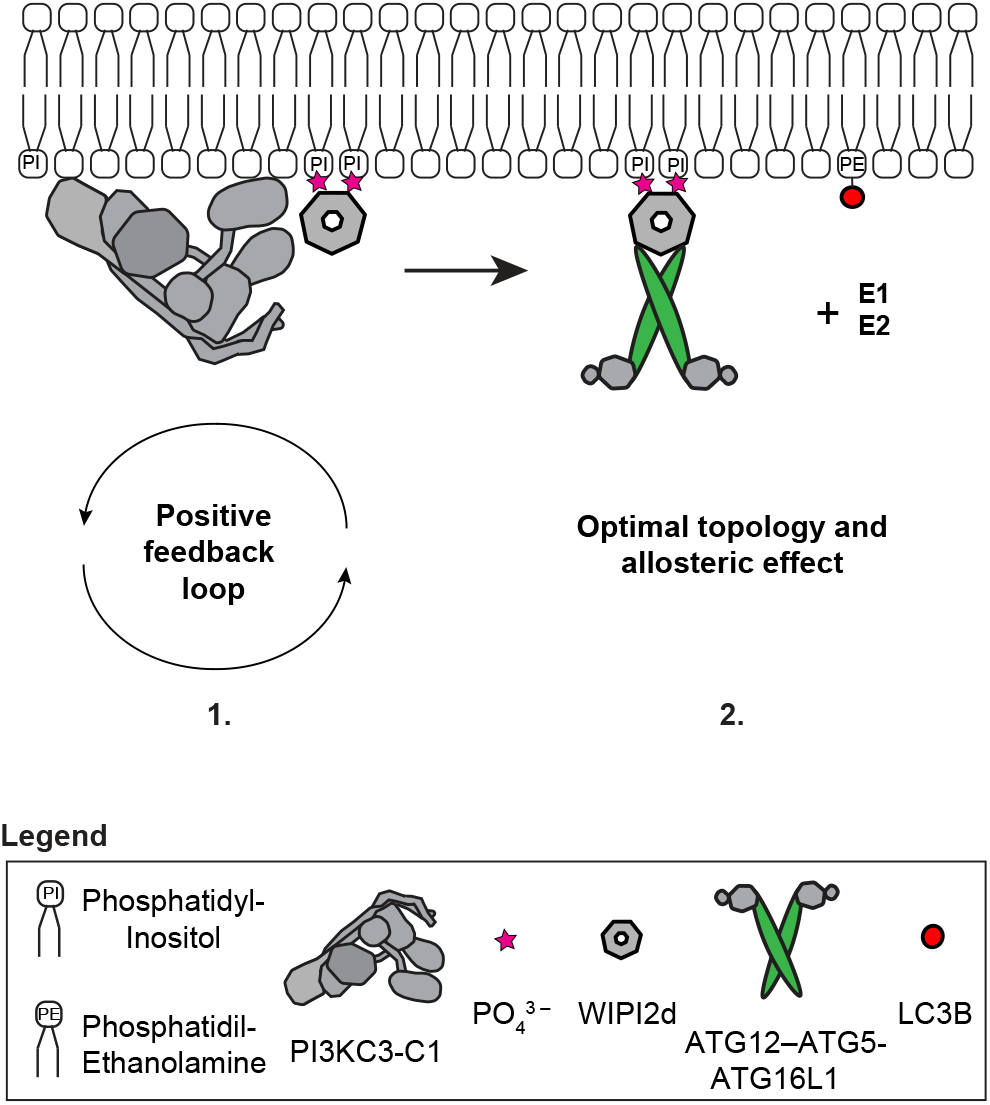
A model of the biochemical reactions reconstituted in this work driving LC3 lipidation *in vivo:* 1) the PI3KC3-C1 complex phosphorylates Phosphatidyl-Inositol (PI) to produce Phosphatidyl-Inositol,3-Phosphate (PI(3)P) on the target membrane. This, in turn, robustly recruits PI(3)P-sensor WIPI2 protein in a self-enhanced positive feedback loop and 2) leads to the downstream recruitment of the E3-like ligase ATG12-ATG5-ATG16L1 complex. These direct protein-protein interactions sustain and promote the catalytic activity of the E3-like ligase enzyme possibly via an induced conformational change within the E3 or the achievement of an optimal topology of the entire lipidation machinery on the target membrane, resulting in the efficient LC3B-PE conjugation.

The inherent membrane binding activity of the human E3 (Dudley et al., 2019; Lystad et al., 2019) is insufficient to recruit it to flat membranes. Here, we confirmed previous observations (Dooley et al., 2014) that WIPI2 is essential for functional E3 recruitment onto membranes resembling flat portions of the ER membrane. Moreover, we discovered that WIPI2d does not merely target E3 to membranes, but also potently activates the LC3 conjugation reaction catalyzed by E3. This effect would be most readily explained by allosteric communication between the WIPI2d binding site on ATG16L1 and the ATG3 binding site on the active ATG12–ATG5 unit (Metlagel et al., 2013; Otomo et al., 2013) of the complex. Alternatively or in addition, WIPI2d might reorient the E3 on the membrane to promote LC3 conjugation (**Figure 6**). It has been suggested that the long-coiled coil of ATG16L1 could span the gap between the omegasome and the phagophore, such that ATG16L1 anchored by omegasome-localized WIPI2 could conjugate LC3 to the phagophore *in trans* (Wilson et al., 2014). This is an attractive model, but our data show that it is at least possible for WIPI2 to promote efficient LC3 lipidation *in cis* on the same membrane. Structural studies will be needed to reveal these mechanisms in more detail.

When these two novel principles, the WIPI2-PI3KC3-C1 positive feedback and the WIPI2 allosteric activation are combined in a single reaction, a massive dose-dependent acceleration of LC3 conjugation is observed. LC3 lipidation nears saturation in ~10 min, which is comparable to what is seen in cells (Axe et al., 2008; Kageyama et al., 2011; Karanasios et al., 2013; Koyama-Honda et al., 2013; Zachari et al., 2019). This is emphasized by the data shown in Figure 4E. These data explain the observation of a PI3KC3-C1 requirement for efficient LC3 lipidation in cells (Dooley et al., 2014)(Axe et al., 2008; Itakura et al., 2008; Itakura and Mizushima, 2010; Matsunaga et al., 2009; Zhong et al., 2009) and of ERGIC-derived membranes in a cell free system (Brier et al., 2019). They also explain the strict requirement for WIPI2 for the lipidation reaction in canonical autophagy (Dooley et al., 2014) and even in STING-induced LC3 lipidation, which bypasses PI3KC3-C1 but is dependent on WIPI2 and the E3 (Gui et al., 2019). These observations place WIPI2 at a truly central position in autophagy initiation and autophagosome biogenesis. These observations also highlight how unexpected properties emerge when multiple steps in autophagy are combined in simultaneous reactions, properties whose quantitative nuances are clarified in the setting of *in vitro* reconstitution. In addition, the data show how combining reactions that each have their own amplification characteristics can drive the overall process with great increased efficiency, helping to explain how autophagosomes are formed *de novo* in cells in a matter of minutes.

## Materials and Methods

Accession numbers: ATG3: NP_071933.2; ATG4: NM_013325.5; ATG5: AGC52703.1; ATG7: NP_001336161.1; mATG7: NP_001240647.1; ATG10: NP_001124500.1; ATG12: O94817.1; ATG16L1: NP_110430.5; LC3B: NP_073729.1; Atg21: KZV07417; WIPI2d: NP_001028691.1; WIPI3: NP_062559.2; WIPI4: NP_001025067.1; ATG14: NP_055739.2; BECN1: NP_003757.1; VPS34: NP_002638.2; VPS15: AAI27106.1; Hrs FYVE: NP_004703.

### Protein expression and purification

Genes coding for protein sequences of human ATG5, ATG12, ATG16L1 (isoform β), ATG7 and ATG10 were codon optimised for Sf9 insect cell expression system and synthetic genes were purchased from GenScript. (10xHis-TEVcs-)ATG16L1-GFP(-TEVcs-StrepII) CDS was assembled via Gibson strategy using the insect codon optimised ATG16L1 gene sequence and monomeric GFP gene sequence. All the ORFs and their tags (see **Table 1**) were inserted into pLIB or pBIG library vectors (Weissmann et al., 2016) via classical restriction cloning. Human ATG12, (10xHis-TEVcs-)ATG5 and human ATG7, ATG10 poli-cystronic constructs were assembled via biGBac system approach (Weissmann et al., 2016) using Gibson assembly strategy. The human ATG12, ATG5, ATG16L1(±GFP), ATG7, ATG10 poli-cystronic gene constructs were cloned via Golden Gate approach by the VBCF Protech Facility (Vienna BioCenter, Vienna).

**Table 1.**
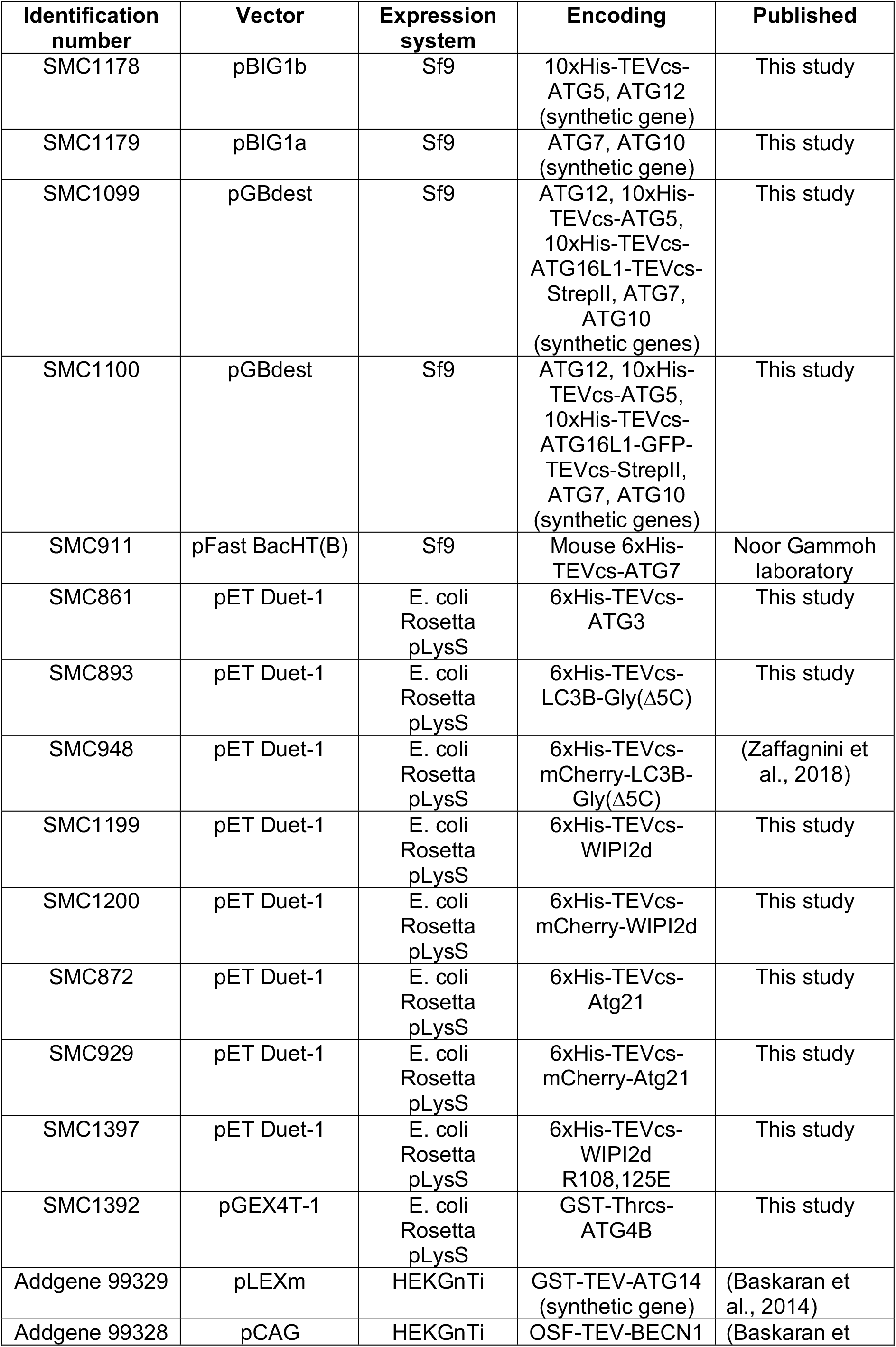
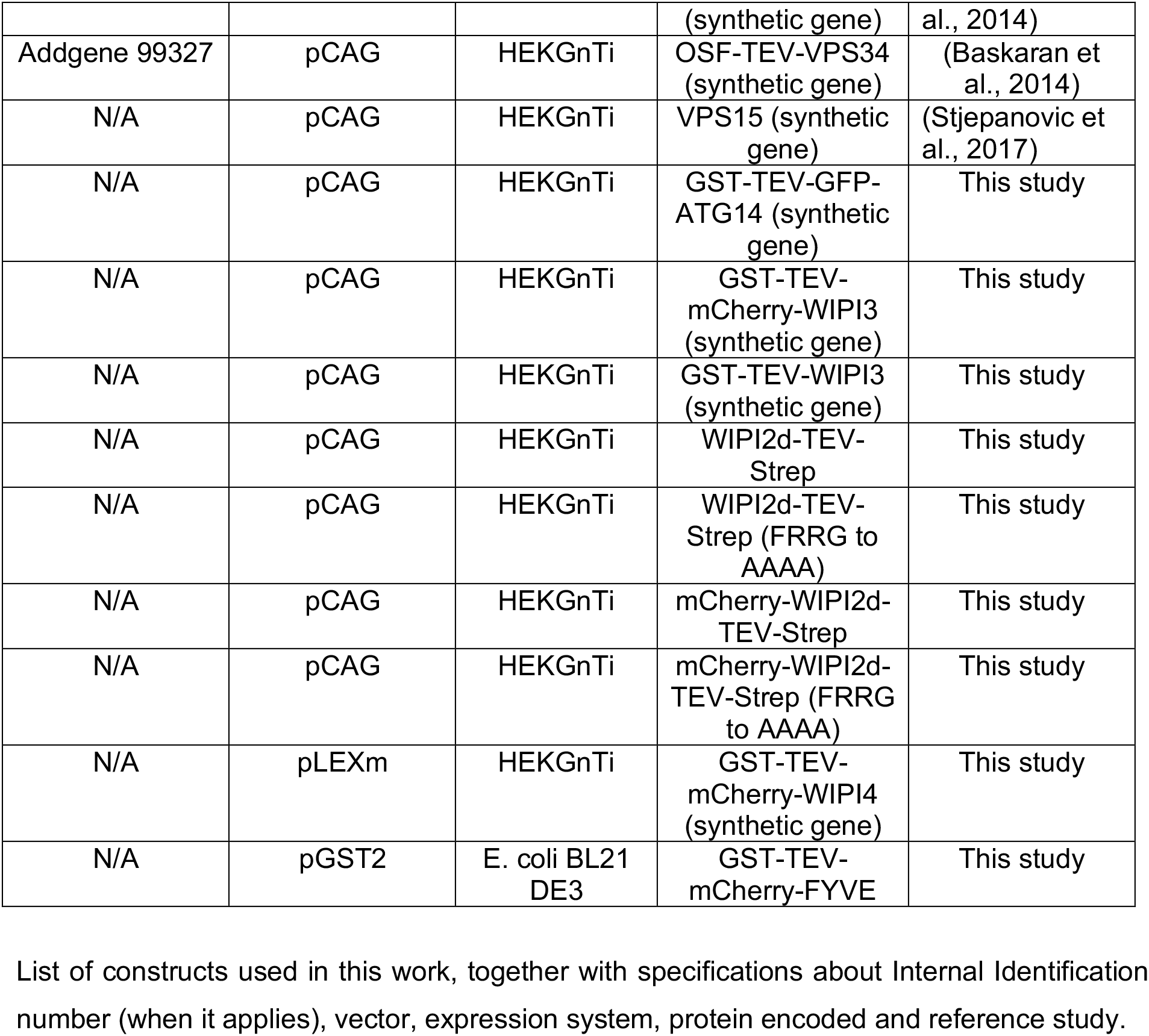
List of constructs used in this work, together with specifications about Internal Identification number (when it applies), vector, expression system, protein encoded and reference study.

2.5 μg of bacmid DNA per construct obtained from amplification in DH10BacY cells were used to transfect 1 Mil Sf9 cells/per construct using Fugene transfection reagent (FuGENE HD, Promega). Virus at P0 was harvested and used to produce a stock Virus P1 solution to further infect 1 lt culture of Sf9 cells at 0.8-1 Mil/ml in SF921 medium containing Pen/Strep. Cultures were harvested when cells reached a viability of max. 95-98%, pelleted down and further washed in 1xPBS at 4000 rpm for 10 minutes at 4°C. Pellets were flash frozen in liquid nitrogen and stored at −80°C until purification.

For expression of (10xHis-TEVcs-)ATG5-ATG12 conjugate, Sf9 cells were co-infected with Virus stocks P1 of the two poli-cystronic constructs coding for ATG12, (10xHis-TEVcs-)ATG5 and ATG7, ATG10, respectively. For purification pellets were thawed and resuspended in ice cold buffer containing 50 mM Hepes pH 7.5, 300 mM NaCl, 10 mM imidazole, 2 mM MgCl_2_, 2 mM β-mercaptoethanol, Complete Protease Inhibitors (Roche), Protease Inhibitor Cocktail (Sigma) and Benzonase Nuclease (Sigma). Cells were lysed on ice by extrusion in a tissue homogenizer and lysates were cleared by ultracentrifugation at 25000 rpm for 45 minutes at 4°C using a Beckman Ti45 rotor. Supernatant was applied to a 5 ml Ni-NTA column (GE Healthcare, Sweden) and eluted via a stepwise imidazole gradient (50, 75, 100, 150, 200, and 300 mM). Protein eluted in fractions containing 150 mM Imidazole. These fractions were pooled, concentrated down and applied onto a Superdex 200 Increase 10/300 GL (GE Healthcare) and eluted in a buffer containing 25 mM HEPES pH 7.5, 150 mM NaCl and 1 mM dithiothreitol (DTT). Fractions containing pure 10xHis-TEVcs-ATG5–ATG12 were pooled, concentrated, snap frozen in liquid nitrogen and stored at −80°C.

For expression of ATG12–(10xHis-TEV-)ATG5-(10xHis-TEVcs-)ATG16L1-(±GFP)-TEVcs-StrepII complexes, Sf9 cells were infected with a single Virus stock P1 corresponding to the poli-cystronic construct coding for GFP-tagged (or not) E3 complex. Cell pellets were thawed and resuspended in ice cold buffer containing 50 mM Hepes pH 7.5, 300 mM NaCl, 2 mM MgCl_2_, 1 mM dithiothreitol (DTT), Complete Protease Inhibitors (Roche), Protease Inhibitor Cocktail (Sigma) and Benzonase Nuclease (Sigma). Cells were lysed on ice by extrusion in a tissue homogenizer and lysates were cleared by ultracentrifugation at 25000 rpm for 45 minutes at 4°C in a Beckman Ti45 rotor. Supernatant was applied to a 5 ml StrepTactin column (GE Healthcare, Sweden) and eluted with 2.5 mM desthiobiotin in 25 mM HEPES pH 7.5, 150 mM NaCl and 1 mM dithiothreitol (DTT). Fractions containing the protein complex were pooled, concentrated down, applied onto a Superdex 6 column (10/300 Increase, GE Healthcare) and eluted in a buffer containing 25 mM HEPES pH 7.5, 300 mM NaCl and 1 mM dithiothreitol (DTT). Fractions containing pure ATG12–(10xHis-TEV-)ATG5-ATG16L1-(±GFP)-StrepII complex were pooled, concentrated, snap frozen in liquid nitrogen and stored at −80°C.

Mouse (6xHis-TEVcs-)ATG7 was expressed in Sf9 cells and harvested following the same procedure used for the ATG12 conjugation machinery constructs described above. For purification, the pellets were treated as for the (10xHis-TEVcs-)ATG5–ATG12 conjugate. Final protein is eluted with a buffer containing 25 mM HEPES pH 7.5, 150 mM NaCl and 1 mM dithiothreitol (DTT), snap frozen in liquid nitrogen and stored at −80°C or −150°C for longer storage.

Human (6xHis-TEVcs-)ATG3 was expressed in E. coli Rosetta pLySS cells. Cells were grown in Luria Bertani (LB) medium at 37°C until an OD600 of 0.4. The culture is than brought to 18°C and grown to an OD600 of 0.8. Protein expression is induced with 100 μM IPTG and grown for a further 16h at 18°C. Cells were pelleted and resuspended in a buffer containing 50 mM HEPES pH 7.5, 300 mM NaCl, 10 mM imidazole, 2 mM MgCl_2_, 2 mM β-mercaptoethanol, 1 mM Pefablock, Roche complete protease inhibitors and DNAse (Sigma). Cells were lysed by freeze thawing and 2×30 s sonication. Lysates were cleared by ultracentrifugation (40000 rpm for 30 min at 4°C in a Beckman Ti45 rotor). Supernatant was filtered (0.45 μm) applied to a 5 ml Ni-NTA column (GE Healthcare, Sweden) and eluted via a stepwise imidazole gradient (50, 75, 100, 150, 200, and 300 mM). Fractions containing the protein of interest were pooled, concentrated down and applied onto a Superdex 75 column (16/60 prep grade, GE Healthcare) and eluted with a buffer containing 25 mM HEPES pH 7.5, 150 mM NaCl and 1 mM dithiothreitol (DTT). Fractions containing pure (6xHis-TEVcs-)ATG3 protein were pooled, concentrated, and stored in final 30% Glycerol concentration at −20°C.

(6xHis-TEV-)LC3BΔ5C and 6xHis-TEV-mCherry-LC3BΔ5C were expressed in *E. coli* Rosetta pLySS cells. Cells were grown in Luria Bertani Broth (LB) medium at 37°C until an OD600 of 0.4. Then, the culture was brought to 18°C and grown to an OD600 of 0.8. Protein expression was induced with 100 μM IPTG and grown for a further 16h at 18°C. Cells were pelleted and resuspended in a buffer containing 50 mM HEPES pH 7.5, 300 mM NaCl, 10 mM imidazole, 2 mM MgCl_2_, 2 mM β-mercaptoethanol, Roche complete protease inhibitors and DNAse (Sigma). Cells were lysed by freeze thawing and 2×30 s sonication. Lysates were cleared by ultracentrifugation (40000 rpm for 30 min at 4°C in a Beckman Ti45 rotor). Supernatant was filtered (0.45 μm) applied to a 5 ml Ni-NTA column (GE Healthcare, Sweden) and eluted via a stepwise imidazole gradient (50, 75, 100, 150, 200, and 300 mM). Fractions containing the proteins of interest were pooled and the 6xHistidine tag was cleaved o.n. at 4°C with TEV protease (only for 6xHis-TEV-LC3B). After cleavage, the sample was concentrated down and applied onto a Superdex 75 column (16/60 prep grade, GE Healthcare) and eluted with a buffer containing 25 mM HEPES pH 7.5, 150 mM NaCl and 1 mM dithiothreitol (DTT). Fractions containing pure 6xHis-TEV-LC3BΔ5C or 6xHis-TEV-mCherry-LC3BΔ5C proteins were pooled, concentrated, snap frozen in liquid nitrogen and stored at −80°C.

WIPI2d gene coding sequence was amplified from a HeLa cDNA library and cloned via classical restriction cloning in a pET-Duet1 vector, with and without a N-terminal mCherry tag, in frame with a N-terminal 6xHistidine tag. The (6xHis-TEV-)WIPI2d R108E,125E mutant construct was obtained by consecutively inserting the two point mutations in the sequence of the wt gene sequence using RTH-PCR strategy. (6xHis-TEV-)WIPI2d wt and R108E,R125E mutant and (6xHis-TEV-)mCherry-WIPI2d were expressed in E. coli Rosetta pLySS cells. Cells were grown in Terrific Broth (TB) medium at 37°C until an OD600 of 0.4. Then, the culture was brought to 18°C and grown to an OD600 of 0.8. Protein expression was induced with 100 μM IPTG and grown for a further 16h at 18°C. Cells were pelleted and resuspended in a buffer containing 50 mM HEPES pH 7.5, 300 mM NaCl, 10 mM imidazole, 2 mM MgCl_2_, 2 mM β-mercaptoethanol, Roche complete protease inhibitors and DNAse (Sigma). Cells were lysed by freeze thawing and 2×30 s sonication. Lysates were cleared by ultracentrifugation (25000 rpm for 30 min at 4°C using a Beckman Ti45 rotor). Supernatant was filtered (0.45 μm) applied to a 5 ml Ni-NTA column (GE Healthcare, Sweden) and eluted via a stepwise imidazole gradient (50, 75, 100, 150, 200, and 300 mM). Fractions at 100-150 mM Imidazole containing the proteins of interest were pooled, concentrated down and applied onto a Superdex 200 column (16/60 prep grade, GE Healthcare) and eluted with a buffer containing 25 mM HEPES pH 7.5, 150 mM NaCl and 1 mM dithiothreitol (DTT). Fractions containing pure protein are pooled, concentrated, snap frozen in liquid nitrogen and stored at −80°C.

*S. cerevisiae* Atg21 gene coding sequence was amplified from a *S. cerevisiae* cDNA library and cloned via classical restriction cloning in a pET-Duet1 vector, with a N-terminal 6xHistidine tag followed by a TEV cleavage (and mCherry tag where it applies) site in frame with the protein coding sequence. (6xHis-TEV-)Atg21 and (6xHis-TEV-)mCherry-Atg21 were expressed in E. coli Rosetta pLySS cells. Cells were grown in Terrific Broth (TB) medium (for Atg21) or LB (Luria Bertani) medium (for mCherry-Atg21) at 37°C until an OD600 of 0.4. Then, the culture was brought to 18°C and grown to an OD600 of 0.8. Protein expression was induced with 100 μM IPTG and grown for a further 16h at 18°C. Cells were pelleted and resuspended in a buffer containing 50 mM HEPES pH 7.5, 300 mM NaCl, 10 mM imidazole, 2 mM MgCl_2_, 2 mM β-mercaptoethanol, Roche complete protease inhibitors and DNAse (Sigma). Cells were lysed by freeze thawing and 2×30 s sonication. Lysates were cleared by ultracentrifugation (40 000 rpm for 30 min at 4°C in a Beckman Ti45 rotor). Supernatant was filtered (0.45 μm) applied to a 5 ml Ni-NTA column (GE Healthcare, Sweden) and eluted via a stepwise imidazole gradient (50, 75, 100, 150, 200, and 300 mM). Fractions at 100-150 mM Imidazole containing the proteins of interest were pooled. For mCherry-Atg21, the 6xHis tag was removed with o.n. cut at 4°C using TEV protease. Consequently, pooled fractions were concentrated down and applied onto a Superdex 200 column (16/60 prep grade, GE Healthcare) and eluted with a buffer containing 25 mM HEPES pH 7.5, 150 mM NaCl and 1 mM dithiothreitol (DTT). Fractions containing pure (±mCherry) Atg21 protein were pooled, concentrated, snap frozen in liquid nitrogen and stored at −80°C.

*H. sapiens* ATG4B gene coding sequence was amplified from a Hela cDNA library and cloned via classical restriction cloning in a pGEX4T1 vector, with a N-terminal GST tag followed by a Thrombin cleavage site in frame with the protein coding sequence. Protein was expressed in E. coli Rosetta pLySS cells. Cells were grown in Luria Bertani (LB) medium at 37°C until an OD600 of 0.4. Then, the culture was brought to 18°C and grown to an OD600 of 0.8. Protein expression was induced with 100 μM IPTG and grown for a further 16h at 18°C. Cells were pelleted and resuspended in a buffer containing 50 mM HEPES pH 7.5, 300 mM NaCl, 2 mM MgCl_2_, 1mM DTT, Roche complete protease inhibitors and DNAse (Sigma). Cells were lysed by freeze thawing and 2×30 s sonication. Lysates were cleared by ultracentrifugation (25 000 rpm for 30 min at 4°C in a Beckman Ti45 rotor). Supernatant was incubated with **GSH beads** (GE Healthcare, Buckinghamshire, UK) and washed with low salt (50 mM HEPES pH 7.5, 300 mM NaCl, 1mM DTT) buffer, followed by high salt (50 mM HEPES pH 7.5, 500 mM NaCl, 1mM DTT) and low salt buffers. Beads were incubated o.n. cut with **Thrombin** protease (SERVA, Heidelberg, Germany) at 4°C. The supernatant containing the eluted and cleaved protein was concentrated down and applied onto a Superdex 75 column (16/60 prep grade, GE Healthcare) and eluted with a buffer containing 25 mM HEPES pH 7.5, 150 mM NaCl and 1 mM dithiothreitol (DTT). Fractions containing pure ATG4B protein were pooled, concentrated, snap frozen in liquid nitrogen and stored at −80°C.

The PI3KC3-C1 complex was expressed and purified from HEK293 GnTi cells as described previously (Chang et al., 2019). ATG14, VPS34, VPS15 and BECN1 constructs were transfected to cells using polyethylenimine (Polysciences). After 60 h expression, cells were harvested and lysed with lysis buffer (50 mM HEPES pH 7.4, 1% Triton X-100, 200 mM NaCl, 1 mM MgCl_2_, 10% glycerol, and 1mM TCEP) supplemented with EDTA free protease inhibitors (Roche). The lysate was clarified by centrifugation (15000 rpm for 1 h at 4 °C) and incubated with glutathione Sepharose 4B (GE Healthcare) for 4 h at 4 °C, applied to a gravity column, and washed extensively with wash buffer (50 mM HEPES pH 8.0, 200 mM NaCl, 1 mM MgCl_2_, and 1mM TCEP). The protein complexes were eluted with wash buffer containing 50 mM reduced glutathione, and treated with TEV protease at 4 °C overnight. TEV-treated complexes were loaded on a Strep-Tactin Sepharose gravity flow column (IBA, GmbH). The complexes were eluted with wash buffer containing 10 mM desthiobiotin (Sigma), and applied to Superose 6 16/50 (GE Healthcare) column equilibrated with gel filtration buffer (20 mM HEPES pH 8.0, 200 mM NaCl, 1 mM MgCl_2_, and 1 mM TCEP). Peak fractions were collected and used immediately for subsequent assays.

WIPI3 and WIPI4 were purified from HEK293 GnTi cells by a similar protocol used for PI3KC3-C1 purification. The TEV-treated proteins were directly applied to a Superdex 200 column (16/60 prep grade, GE Healthcare) equilibrated with gel filtration buffer (20 mM HEPES pH 8.0, 150 mM NaCl, and 1 mM TCEP). Fractions containing pure proteins were pooled, concentrated, snap frozen in liquid nitrogen and stored at −80°C.

### Static light Scattering (SLS)

A sample of 90 μl at 10 μM (1,5 mg/ml) of purified ATG12–ATG5-ATG16L1-GFP was applied to a Superose 6 Increase 10/300 GL column in 25 mM Hepes pH 7.5, 300 mM NaCl, 1 mM DTT. The column was coupled to a Wyatt TREOS II instrument (Wyatt Technology Corporation, Santa Barbara, CA, USA). Data were analyzed using the ASTRA V software (Wyatt).

### Preparation of Giant Unilamellar Vesicles (GUVs)

GUVs were prepared by electroformation (Romanov et al., 2012) or hydrogel-assisted swelling (Weinberger et al., 2013), as described previously. To prepare GUVs by electroformation, lipids were desiccated for 5 hrs and the electroformation was conducted at 30°C in a 309 mOsm Sucrose solution. GUVs were used for experiments directly afterwards. To prepare GUVs by gel-assisted swelling, PVA with a molecular weight of 145,000 (Millipore) was used as hydrogel substrate. PVA was mixed with water to obtain a 5% (w/w) PVA solution and stirred on a heat plate at around 90°C until the solution is clear. 300 μl of 5% PVA were then spun coat for 30 s at a speed of 1200 rpm on a plasma-cleaned cover glass with 25 mm diameter. To dry the PVA film, the coated glass was placed for 30 min in a heating incubator at 60°C. 10-15 μl of lipids with different compositions at 1 mg/ml stock were deposited uniformly on the PVA film. Details about different lipid compositions are indicated in Figure legends. The lipid-coated cover glass was put under vacuum overnight to evaporate the solvent from the dissolved lipid mixture. The coated cover glass was transferred into a 30 mm dish and 300 μl 400 mOsm sucrose solution was added on top of the glass. After swelling for around 1 h at room temperature, the vesicles were harvested and then stored at room temperature and used immediately.

Atto647N DOPE (Atto TEC, AD 647N-161, 1 mg/ml) was used as a GUV membrane dye. All the other lipids for GUVs preparation are from Avanti Polar Lipids (Alabaster, AL, USA). GUVs with DO lipids contained: DOPC (850375C, 10 mg/ml), DOPE (850725C, 10 mg/ml), DOPS (840035C, 10 mg/ml), POPI (850142P, 1 mg/ml). GUVs with PO lipids contained: POPC (850457C, 10 mg/ml), POPE (850757C, 10 mg/ml), POPS (840034C, 10 mg/ml), POPI. GUVs with Brain lipids contained: Brain PC (840053C, 10 mg/ml), Brain PE (840022C, 10 mg/ml), Brain PS (840032C, 10 mg/ml), Liver PI (840042C, 10 mg/ml). PI(3)P (850150P, 1 mg/ml) was used for GUVs containing PI(3)P.

### Preparation of Small Unilamellar Vesicles (SUVs)

For preparation of Small Unilamellar Vesicles (SUVs) lipids were mixed to homogeneity in Chloroform and dried under a stream of Argon and further desiccated for one hour under vacuum. The lipid film was rehydrated in 25 mM HEPES pH 7.5, 137 mM NaCl, 2.7 mM KCl,0.1 mM DTT. The lipid film was resuspended by gentle mixing at RT and sonicated for 2 min in a bath sonicator. The resuspended SUVs were then extruded 21 times through a 0.4 μm membrane followed by 21 times through a 0.1 μm membrane (Whatman, Nucleopore, UK) using the Mini Extruder from Avanti Polar Lipids Inc.. The final SUV suspension has a concentration of 1 mg lipids/ml.

Lipids used for SUVs preparation are from Avanti Polar Lipids: DOPC (850375C, 10 mg/ml), DOPE (850725C, 10 mg/ml). Alternatively, when PO-fatty acid chains were used SUVs contained: POPC (850457C, 10 mg/ml), POPE (850757C, 10 mg/ml). Other lipids: ATTO390-PE ((ATTO-TEC, 1mg/ml), PI(3)P (850150P, 1 mg/ml) prepared as in (Fracchiolla et al., 2016) and 15% liver PI (840042C, 10 mg/ml).

### Liposome sedimentation assay

SUVs prepared as described in resuspended in **Preparation of Small Unilamellar Vesicles (SUVs)** were mixed with 1-2 μg protein mixes at a ratio of 1:1 in buffer containing 25 mM HEPES pH 7.5, 137 mM NaCl, 2.7 mM KCl, 0.1 mM DTT. Reactions were incubated with 0.5 mg/ml liposomes for 30 min at room temperature (22°C). Then the reactions were centrifuged at 180000 g at 22°C, supernatants (S) and pellets (P) were separated and equal amounts were run on 4-15% SDS/polyacrylamide gels and stained with Coomassie Brilliant Blue.

### SUVs membrane recruitment assay on beads

For experiments shown in **Figure S3B, C** GFP-Trap (**Figure S3B**) or RFP-Trap (**Figure S3C**) beads (Chromotek) were first coated with the ATG12–ATG5-ATG16L1-GFP protein (**Figure S3B**) or mCherry or mCherry-WIPI2d or mCherry-Atg21 proteins (**Figure S3C**) and where it applies, upon washings with buffer containing 25 mM HEPES pH 7.5, 137 mM NaCl, 2.7 mM KCl, 0.1 mM DTT further incubated with mCherry-tagged Proppins (**Figure S3B**). After washings, 1-2 μl of beads per sample were pipetted into the wells of a 384-well glass-bottom microplate (Greiner Bio-One) pre-filled with a prep of ATTO390-SUVs in buffer containing 25 mM HEPES pH 7.5, 137 mM NaCl, 2.7 mM KCl, 0.1 mM DTT at a concentration of 0,2 mg/ml.

### Membrane protein recruitment – GUV assay

For experiment shown in **Figure 1B**, 15 μl of the ATG12–ATG5-ATG16L1-GFP protein are added to 15 μl of GUVs pre-pipetted into the wells of a 384-well glass-bottom microplate (Greiner Bio-One) pre-coated with a 5 mg/ml BSA solution in buffer containing 25 mM Hepes pH=7.5 and 150 mM NaCl, for a final concentration of 500 nM in buffer containing 25 mM HEPES pH 7.5, 137 mM NaCl, 2.7 mM KCl, 0.1 mM DTT. For experiment shown in **Figure 2B**, 15 μl of the ATG12–ATG5-ATG16L1-GFP at 500 nM and mCherry-WIPI2d at 500 nM were added to 15 μl GUVs. Concentrations are calculated for a final volume of 30 μl. The images were acquired after 30 minutes of incubation at room temperature (22°C) in the dark using a LSM700 confocal microscope (Zeiss) with a 20x/0.8 Plan Apochromat objective (lasers 488 nm, 10mW - 555 nm, 10mW), controlled by Zeiss ZEN 2012 Software and processed with ImageJ software. Identical laser power and gain settings were used during the acquisition of all conditions. For experiment shown in **Figure S3D**, 15 μl of the ATG12– ATG5-ATG16L1-GFP at 100 nM and WIPI2d at 200 nM (or buffer) were added to 15 μl GUVs. Concentrations are calculated for a final volume of 30 μl. The images were acquired after at least 30 minutes of incubation at room temperature (22°C) in the dark using a Spinning Disc microscope (Visitron) with a 63x objective and processed with ImageJ software. Identical laser power and gain settings were used during the acquisition of all conditions.

### *In vitro* bulk LC3B lipidation assay

Reactions were set up by mixing 1:1 volume of SUVs and proteins’mix. Protein mix was prepared in buffer containing 25 mM HEPES pH 7.5, 137 mM NaCl, 2.7 mM KCl, 0.1 mM DTT at a final concentration of 1 μM of mouse ATG7, 1 μM of human ATG3, 5 μM of LC3BΔ5C and 1 μM of E3-like ligase or ATG12–ATG5 conjugate (**Figure 1A**). 0.1 μM of E3-like ligase was used in experiments shown in **Figure 2F** together with 0.5 μM of WIPI2d. 0.5 μM of E3-like ligase was used in experiments shown in **Figure 3E, H** together with 2.5 μM of Atg21 or WIPI2d wt or mut. Reactions were conducted on a Thermoblock at 37°C in the presence of 5 mM ATP and 1 mM MgCl_2_. 15 μl of reaction were sampled at each timepoint, mixed with 3 μl of 6x Protein Loading dye, boiled at 60°C for 10 minutes and loaded on SDS PAGE followed by staining with Coomassie Brilliant Blue.

### *In vitro* reconstitution of LC3B lipidation on GUVs

For experiments shown in **Figure 1C** proteins were mixed in 15 μl and added to 15 μl of GUVs pre-pipetted into the wells of a 384-well glass-bottom microplate (Greiner Bio-One), for a final concentration of 100 nM of mouse ATG7, 100 nM of human ATG3, 500 nM E3-GFP and 500 nM of mCherry-LC3BΔ5C, together with 0.5 mM ATP, 0.5 mM MgCl_2_. The reaction buffer contains 25 mM HEPES at pH 7.5, 150 mM NaCl and 0.1 mM DTT.

For experiments shown in **Figure 2C** proteins were mixed in 15 μl and added to 15 μl of GUVs pre-pipetted into the wells of a 384-well glass-bottom microplate (Greiner Bio-One), for a final concentration of 100 nM of mouse ATG7, 100 nM of human ATG3, 100nM E3-GFP and 500 nM of mCherry-LC3BΔ5C, together with 0.5 mM ATP, 0.5 mM MgCl_2_.

Concentrations of proteins and cofactors used are calculated for a final volume of 30 μl reaction. The images were acquired after 30 minutes of incubation at room temperature (22°C) in the dark (**Figure 2C**) using a LSM700 confocal microscope (Zeiss) with a 20x/0.8 Plan Apochromat objective (lasers 488 nm, 10mW - 555 nm, 10mW), controlled by Zeiss ZEN 2012 Software and processed with ImageJ software.

For de-lipidation experiments, 10U of CIP were added to the well together with ATG4B at a final concentration of 1 μM, or buffer in the negative control and GUVs imaged under a Spinning Disc microscope (Visitron) equipped with a 63x Objective and a EM-CCD Camera. For experiments with PI3KC3-C1, the reactions were performed at room temperature in an observation chamber (Lab Tek) coated with 5 mg/ml beta casein. 15-20 μl GUVs were added at last to initiate the reaction with a final volume of 150 μl. The reaction buffer contains 20 mM HEPES at pH 8.0, 190 mM NaCl and 1 mM TCEP. A final concentration of 100 nM E3-GFP, 100 nM ATG7, 100 nM ATG3, and 500 nM mCherry-LC3BΔ5C was used for all lipidation reactions with PI3KC3-C1. The final concentrations of PI3KC3-C1, WIPI2d, or FYVE domain used in different experiments are indicated in figure legends. After 5 min incubation, during which we picked random views for imaging, time-lapse images were acquired in multitracking mode on a Nikon A1 confocal microscope with a 63 × Plan Apochromat 1.4 NA objective. Identical laser power and gain settings were used during the course of all conditions.

### Microscopy-based bead protein-protein interaction assay

mCherry, mCherry-Atg21 and mCherry-WIPI2d bait proteins are incubated at a concentration of 40 μM each with 10 μl RFP-Trap beads (Chromotek) in buffer containing 25 mM HEPES at pH 7.5, 150 mM NaCl and 1 mM DTT (**Figure 3D**) or GFP-Trap beads (Chromotek) when E3-GFP was used (**Figure 2A**). Beads are incubated with proteins for at least 30 minutes at 4°C on a rotating wheel and washed 3 times in 100 μl buffer. 1 μl beads taken from a 1:1 volume beads:buffer mix, are pipetted onto the well of a 384-well glassbottom microplate (Greiner Bio-One) pre-coated with a 5 mg/ml BSA solution in buffer containing 25 mM Hepes pH=7.5 and 150 mM NaCl, containing 15 μl of the ATG12–ATG5-ATG16L1-GFP pray protein at 5 μM. Beads were incubated for at least 30 minutes at room temperature (22°C) and imaged using a LSM700 confocal microscope (Zeiss) (lasers 488 nm, 10mW - 555 nm, 10mW), controlled by Zeiss ZEN 2012 Software with a 20x/0.8 Plan Apochromat objective and processed with ImageJ software.

For interaction between E3 and WIPI3 or WIPI4, a mixture of 500 nM GST-mCherry-WIPI3 or GST-mCherry-WIPI4 and 100 nM E3-GFP is incubated with 10 μl Glutathione Coated Polystyrene Particles (Spherotech) in reaction buffer containing 25 mM HEPES at pH 7.5, 150 mM NaCl and 1 mM TCEP. After incubation at room temperature for 30 min, the beads were washed three times, suspended with 120 μl reaction buffer, and then transferred to the observation chamber for imaging. Images were acquired on a Nikon A1 confocal microscope with a 63 × Plan Apochromat 1.4 NA objective.

### FRAP (Fluorescence Recovery After Photobleaching)

The surface of GUVs treated for LC3B lipidation was photobleached using a Spinning Disc microscope (Visitron) with a 100% laser intensity and 10sec/pixel.

### Western Blotting and Antibodies

Protein samples analysed in **Figure S3D** were subjected to electrophoretic run and Western Blot onto a Nitrocellulose membrane in buffer containing 20% Et-OH. After membrane blocking for 30 min at RT in 3% milk/TBST (0.1% Tween20), the membrane was incubated with mouse anti-WIPI2b primary antibody (BioRad, MCA5780GA) used at a dilution of 1:1000 in 3% milk/TBST (0.1% Tween20). HRP-conjugated goat anti-Mouse (Jackson ImmunoResearch, # 115-035-003, respectively) was used at 1:10000 dilution in 3% milk/TBST (0.1% Tween20). Signal was developed with Clarity ECL substrate (BioRad). The signal was recorded with a ChemiDOC Touch (Biorad) imager.

### Quantifications and statistical analysis

#### GUV image quantification

Image J was used for the data analysis of GUV fluorescence intensities over time. The three channel GUV movies were split to individual channels. The outline of individual vesicles was manually defined based on the membrane fluorescence channel. For each vesicle, the intensity thresholding was calculated by the average intensities of pixels inside and outside of the vesicle. The intensity trajectories frame by frame of individual GUV were then obtained. Multiple intensity trajectories were calculated from multiple data sets and the average and standard deviation calculated and reported.

For quantification shown in **Figure S3D** multiple lines were manually drawn across the GUV membranes identified in the Bright Field channel. GFP signal intensity was measured at these positions as the Max Intensity signal along the line. Averages and Standard deviations were calculated among the measured values per each condition and plotted in a bar graph.

For experiments shown in **Figure 2E, F** multiple lines were manually drawn across the GUV membranes identified in the GFP channel at time 0. The position of the line was checked throughout the time course to cross the GUV membrane. GFP and mCherry signal intensities were simultaneously measured at these positions as the Max Intensity signal along the line. A GUV free area at time point 0 was chosen to measure the background value for the mCherry channel as Mean value. This was substracted from the mCherryintensities measured for all the GUVs of the field of view along the time course. mCherry and GFP signal at time 0 were set to 100% and the corresponding valued of the timecourse were calculated relative to this. Data were plotted ina scatter graph as a function of time.

For FRAP experiments in **Figure 2D**, a line was manually drawn across the GUV membrane within the area subjected to photobleaching. mCherry Max Intensity signal along the line was measured. The value corresponding to the pre-bleached timepoint was set as 100% and all the values measured after photobleaching were related to this and plotted in a scatter graph as function of time. The p-values were calculated using an unpaired two-tailed Student’s *t*-test. P-values were considered as follows: p≥0.5, not significant (ns); 0.01<p<0.05 (*); 0.001<p<0.01 (**); 0.0001<p<0.001 (***); p<0.001 (****).

#### Bead images quantification

Image J software was used for beads images quantifications. Multiple lines were manually drawn across the beads. These lines were saved as Regions Of Interest (ROI). mCherry and GFP signal intensity were simultaneously measured at these ROIs as the Max Intensity signal along the line. Averages and Standard deviations were calculated among the measured values per each condition and plotted in a bar graph.

#### SUV sedimentation

The gels of three independent experiments were quantified using the Analyze Gel tool of Image J software. The amounts of pelleted protein per each condition were measured as the area below the corresponding peak. The measured amount of pelleted protein in the absence of SUVs was subtracted from the measured amount of pelleted protein in the presence of SUVs. These values were plotted as relative or absolute pelleted protein amounts in bar graphs.

#### In vitro LC3B lipidation on SUVs

The gels of three independent experiments were quantified using the Analyze Gel tool of Image J software. The amounts of LC3B-I and -II at each timepoint were measured as the area below the corresponding peak. The sum of the values corresponding to LC3B-I and -II peaks was then calculated at each timepoint and the fraction of LC3B-II was calculated. Finally, the relative LC3B-II fraction amounts were calculated at each timepoint and plotted as a function of time.

When a contaminant band of WIPI2d batch with a running behaviour similar to that of LC3B-II was present and overlapped with the protein of interest (**Figure 3D**) the value corresponding to the contaminant protein peak present at timepoint 0 was subtracted from each value measured for the LC3B-II band in the following timepoints. Instead, in Figure 3F the value assigned to LC3B-II was arbitrarily set to 0, as both wt and mutant proteins had the same contaminant band, not be present at the following timepoint. The analysis followed as described above.

## Acknowledgements

This work was supported by Human Frontiers Science Program RGP0026/2017 (J.H.H. and S.M.), ERC grant No.646653 (S.M.) and NIH R01 GM111730 (J.H.H.). We thank the Max Perutz Labs Mass Spectrometry, Max Perutz Labs BioOptics and VBCF Protech Facilities for technical support and the VBCF for providing the MS instrument pool. We thank Noor Gammoh for sharing the plasmid encoding mouse ATG7.

## Conflict of interest

J.H.H. is a co-founder of Casma Therapeutics. S.M is member of the scientific advisory board of Casma Therapeutics.

## Author contributions

D. Fracchiolla, C. Chang, J.H. Hurley, and S. Martens designed all experiments. D. Fracchiolla, and C. Chang performed experiments and prepared the figures. D. Fracchiolla, C. Chang, J.H. Hurley, and S. Martens wrote the manuscript.

## Abbreviations

(E3): ATG12–ATG5-ATG16L1
(ATG): Autophagy
(PI3KC3-C1): Class III phosphatidylinositol-3 kinase complex I
(DO): dioleoyl
(GUV): Giant Unilamellar Vesicle
(LC3B): Microtubule-associated proteins 1A/1B light chain 3B
(PO): palmitoyl-oleoyl
(SUV): Small Unilamellar Vesicle
WIPI: (WD-repeat protein interacting with phosphoinositides)

## Supplemental Material

The Supplemental Material section contains Supplemental Figures 1-4 including their legends, and a summary table **(Table 1)** listing all the constructs used in this work.

**Figure S1.**
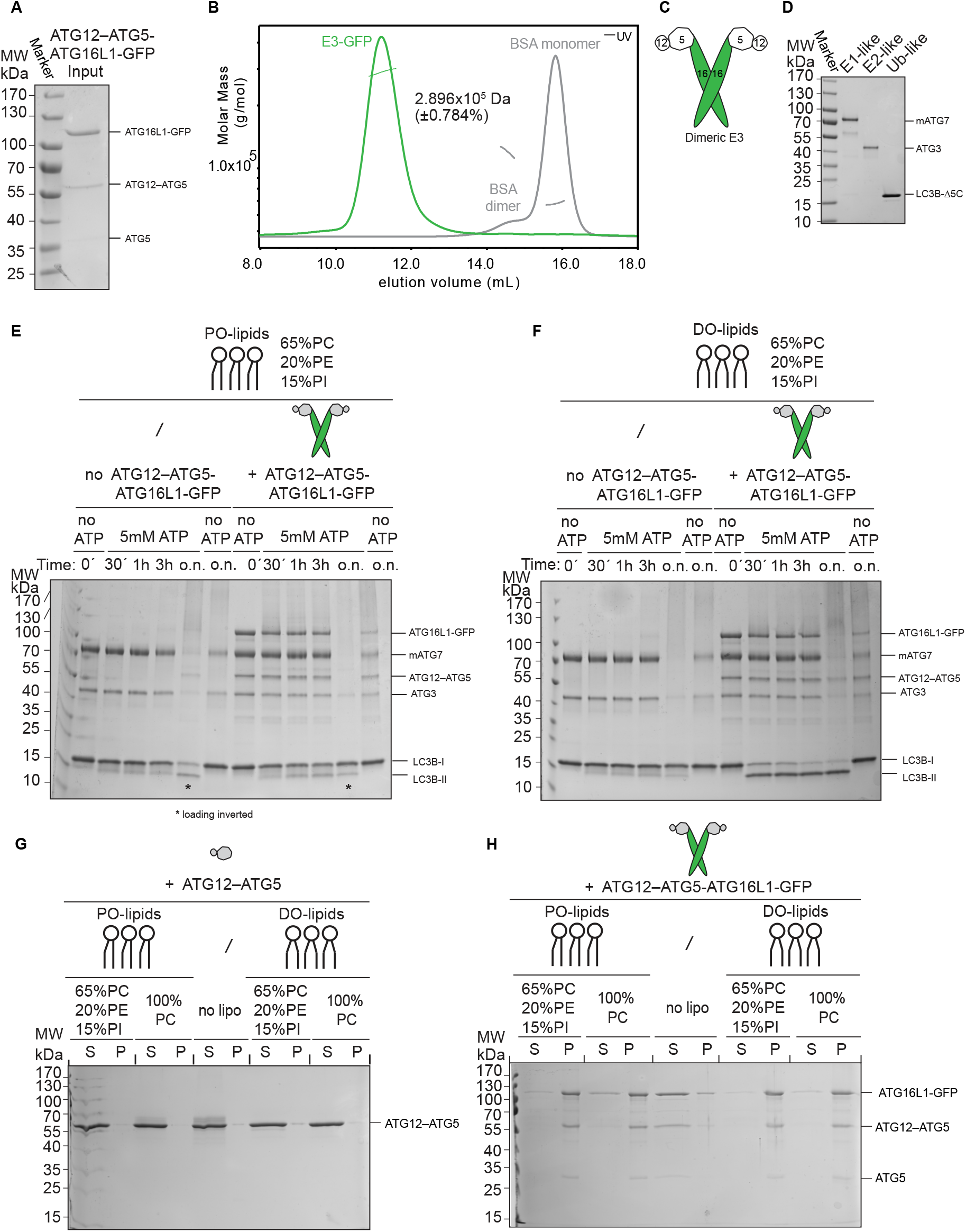
Biochemical characterization of the E3 and the LC3 lipidation machinery. **A** Purified human E3-GFP complex resolved on a 10% SDS PAGE and stained with Coomassie Brilliant Blue. **B** Static Light Scattering (SLS) plot of recombinant E3-GFP. The protein was applied onto a Superose 6 Increase 10/300 GL column coupled with a Wyatt TREOS II instrument. BSA was used for calibration. **C** Schematic representation of the dimeric E3-like ligase holo-complex containing two copies of each subunit of ATG12, ATG5 and ATG16L1(+monoGFP). **D** Recombinant mouse ATG7, human ATG3 and LC3BΔ5C resolved on a 10% SDS PAGE and stained with Coomassie Brilliant Blue. **E** *In vitro* LC3B lipidation assay employing PO-SUVs (65% PC: 15% liver PI: 20% PE), ATG7, ATG3, E3-GFP (1 μM) and LC3B incubated at 37°C in the presence of MgCl_2_/ATP. Samples taken at the indicated time points (o.n.: overnight) were loaded on a 4-15% SDS PAGE. Timepoints corresponding to o.n. incubation indicated with asterisk (*) were swapped during loading. **F** ATG7, ATG3, E3-GFP (1 μM) and LC3B were incubated with DO-SUVs (65% PC: 15% PI: 20% PE) at 37°C with MgCl_2_ and ATP. Samples at the indicated time points (o.n.: overnight) were loaded on a 4-15% SDS PAGE. **G** Co-sedimentation assay of ATG12-ATG5 with DO-lipid (left) or PO-lipid (right) SUVs (65% PC: 20% PE: 15% PI). **H** Co-sedimentation assay of the E3 with DO-lipid (left) or PO-lipid (right) SUVs with the indicated lipid composition.

**Figure S2.**
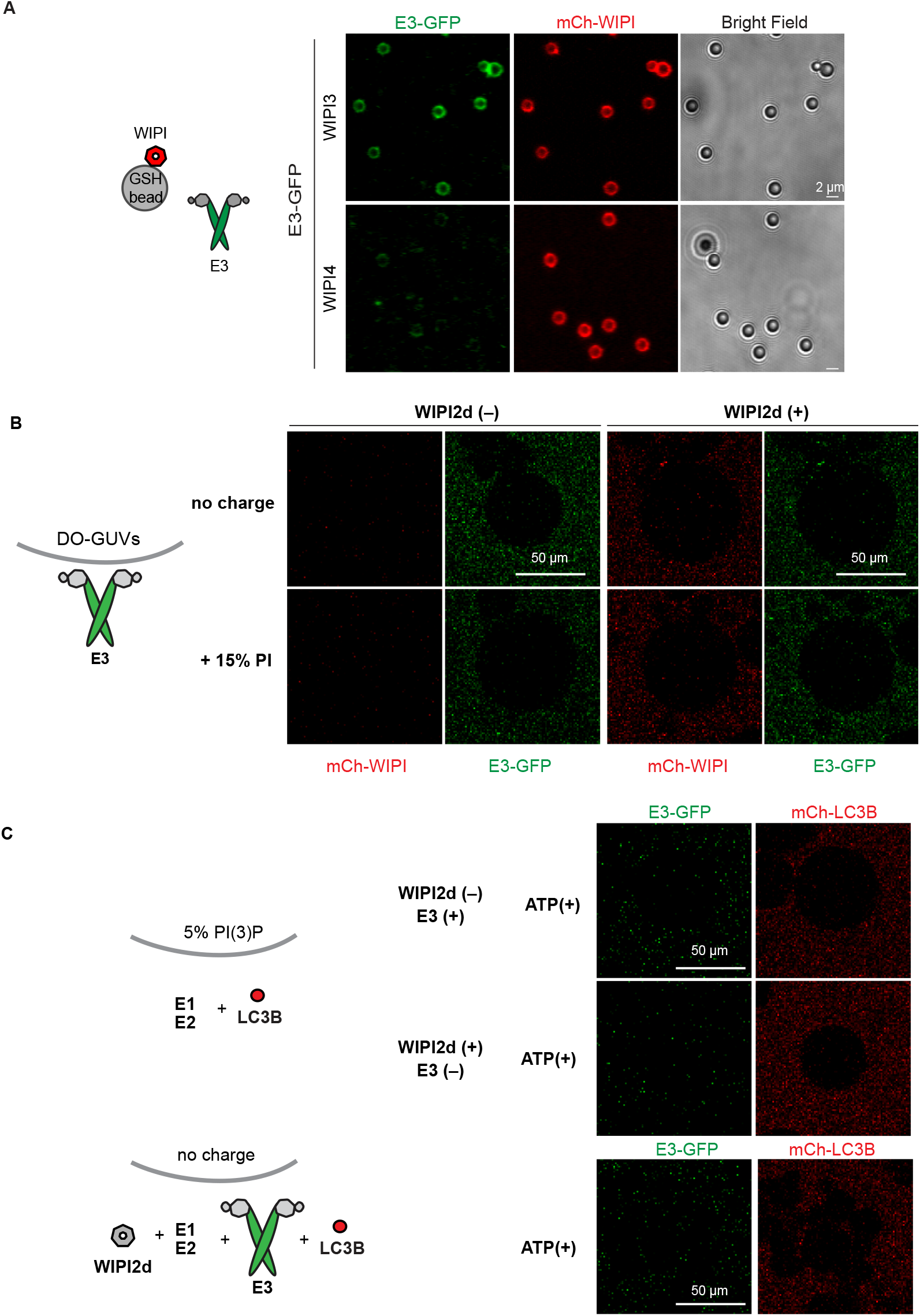
PI(3)P and WIPI2d-dependent E3 recruitment to GUVs and LC3 lipidation. **A** GST-mCherry-tagged WIPI3 or WIPI4 was incubated with GFP-tagged E3 and glutathione coated polystyrene beads. Representative confocal images taken after 30 min incubation are shown. Scale bars, 2 μm. **B** The E3 was co-incubated with mCherry-WIPI2d and DO-GUVs containing either 65% PC: 20% PE: 15% PI (bearing the same net negative charge as GUVs in **Figure 2B)** or GUVs containing 80% PC: 20% PE (zero net charge). Scale bars, 50 μm. **C** mCherry-LC3B was co-incubated with ATG7 and ATG3 in the presence or absence of E3-GFP or WIPI2d with GUVs containing either 65% PC: 20% PE: 15% PI (with the same net negative charge as GUVs in **Figure 2C**) or GUVs containing 80% PC: 20% PE (zero net charge). Scale bars, 50 μm.

**Figure S3A.**
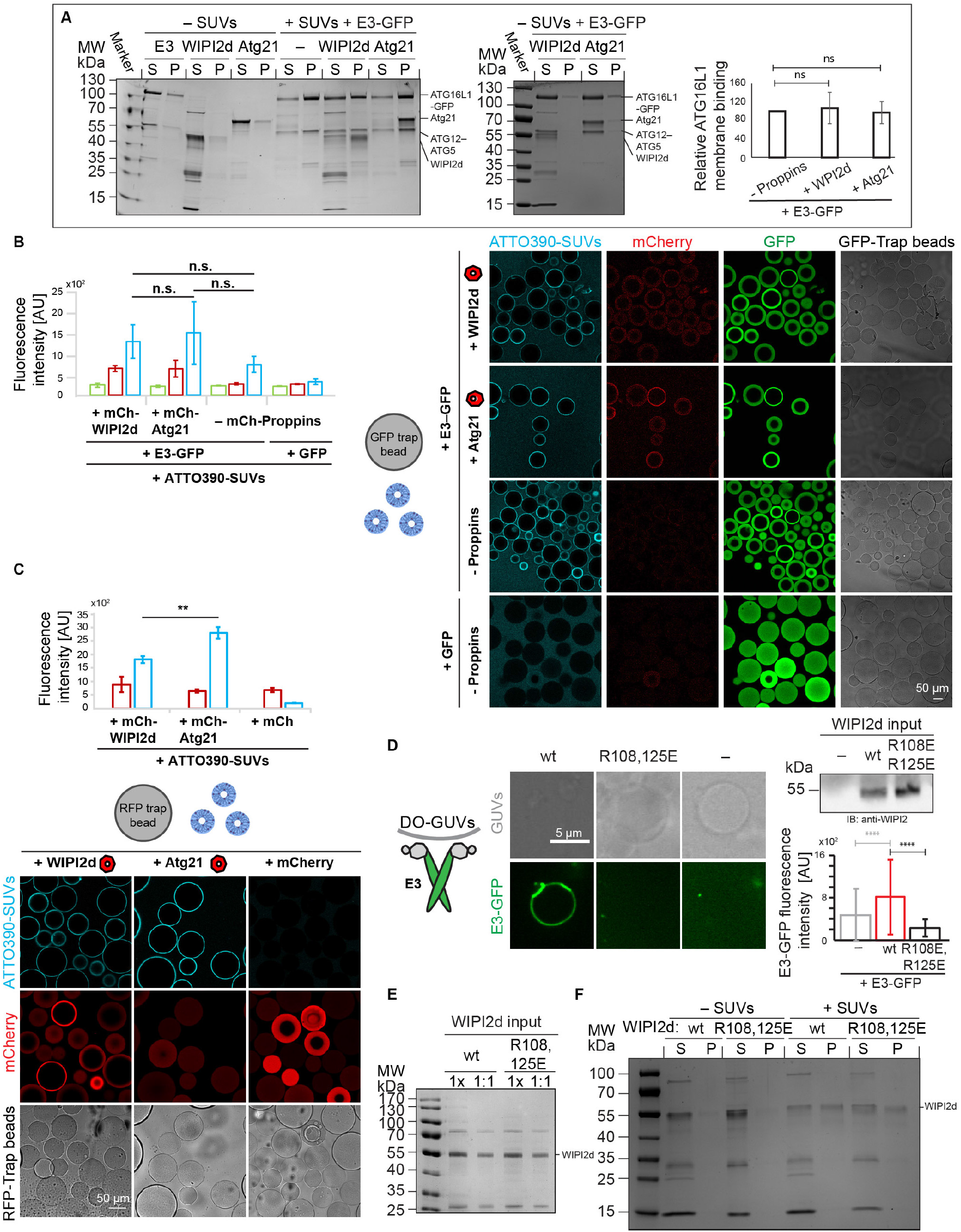
The role of WIPI2d binding to the E3 complex goes beyond its membrane recruitment. **A** Co-sedimentation assay using DO-SUVs (75% PC: 20% PE: 5% PI(3)P) and the E3-GFP complex in the presence or absence of WIPI2d or Atg21. Quantification of three independent experiments (means ± SDs are shown; N = 3) is shown on the right. **B** GFP-Trap beads coated with E3-GFP and mCherry-WIPI2d or mCherry-Atg21 or no PROPPINs were incubated with DO-SUVs containing 72% PC: 20% PE: 5% PI(3)P, 3% ATTO390-PE. Quantification of three independent experiments (means ± SDs are shown; N = 3) is shown. **C** RFP-Trap beads coated with mCherry, mCherry-WIPI2d or mCherry-Atg21 were incubated with DO-SUVs containing 72% PC: 20% PE: 5% PI(3)P, 3% ATTO390-PE. Quantification of three independent experiments (means ± SDs are shown; N = 3) is shown. **D** E3-GFP (0.1 μM) recruitment to DO-GUVs containing 75% PC: 20% PE: 5% PI(3)P in the presence of wt WIPI2d (0.2 μM) or R108,125E mut WIPI2d (0.2 μM) or no PROPPINs. Scale bar 5 μm. The E3-GFP signal on GUVs was quantified and plotted (means ± SDs are shown; N wt= 107, N mut= 197, N no PROPPINs= 170). A blot probed for WIPI2 shows the protein input. **E** Coomassie stained gel showing equal amounts of wt and R108,125E mutant proteins used in the bulk lipidation assays of **Figure 3G, H.** **F** Co-sedimentation assay using DO-SUVs (75% PC: 5% PI(3)P: 20% PE) and the wt WIPI2d or R108,125E WIPI2d. P-values were calculated using Student’s *f*-test (p≥0.5 (ns); 0.01<p<0.05 (*); 0.001<p<0.01 (**); 0.0001<p<0.001: (***); p<0.001: (****)).

**Figure S4.**
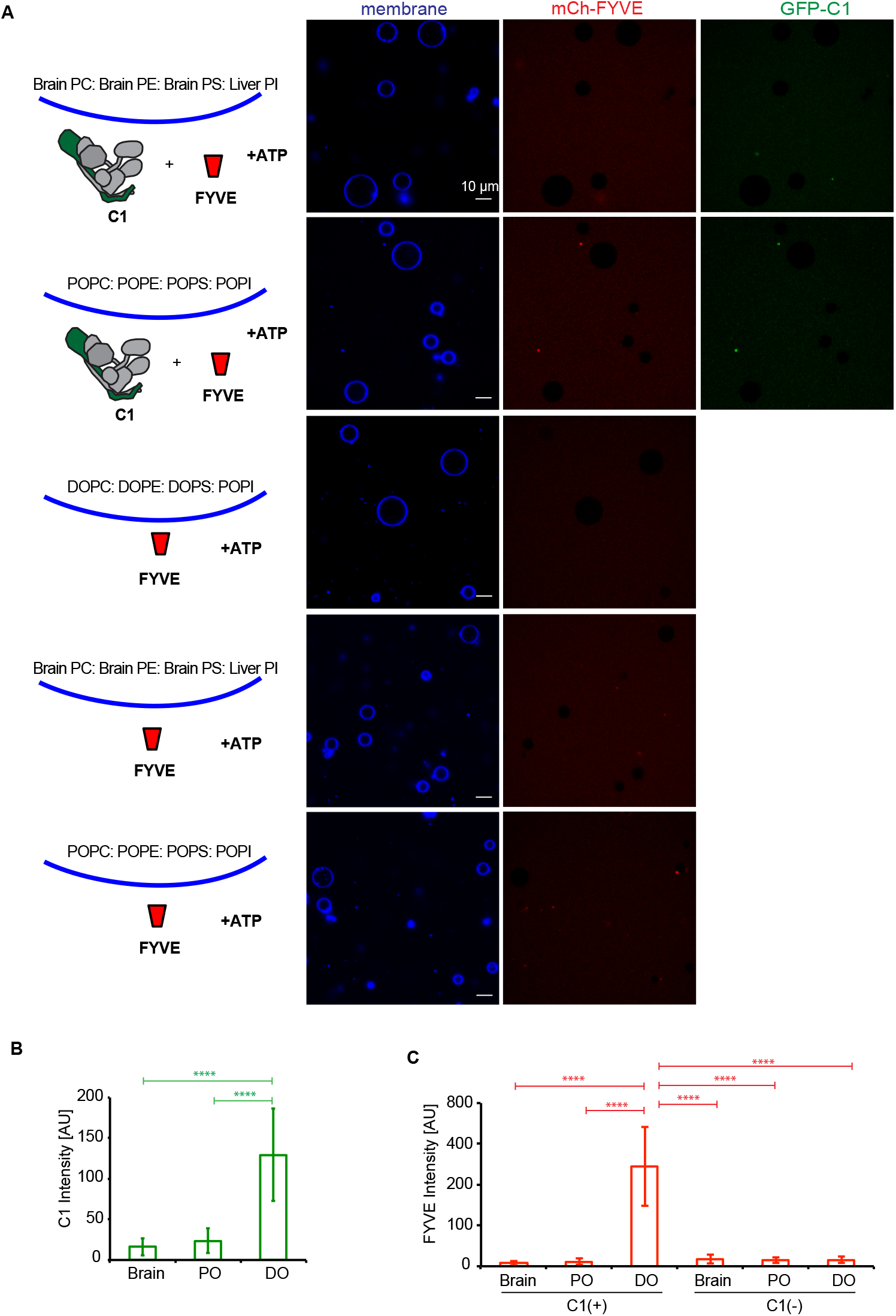
PI3KC3-C1 is active on unsaturated flat membranes. **A** Representative confocal images of GUVs showing the binding of the PI3KC3-C1 complex and FYVE domain on different membranes. mCherry-FYVE domain (1 μM) was incubated with GUVs with brain lipids (64.8% Brain PC: 20% Brain PE: 5% Brain PS: 10% liver PI:0.2% Atto647 DOPE), PO lipids (64.8% POPC: 20% POPE: 5% POPS: 10% POPI: 0.2% Atto647 DOPE), or DO lipids (64.8% DOPC: 20% DOPE: 5% DOPS: 10% POPI: 0.2% Atto647 DOPE) in the absence or presence of GFP-tagged PI3KC3-C1 (200 nM) for 30 min. Scale bars, 10 μm. **B** Quantification of the relative intensities of PI3KC3-C1 on different GUV membranes (means ± SDs; N = 50). P-values were calculated using Student’s *f*-test (p≥0.5: (ns); 0.01<p<0.05: (*)). **C** Quantification of the relative intensities of FYVE domain on different GUV membranes (means ± SDs; N = 50). P-values were calculated using Student’s *f*-test (p<0.001: (****)).

**Figure S5.**
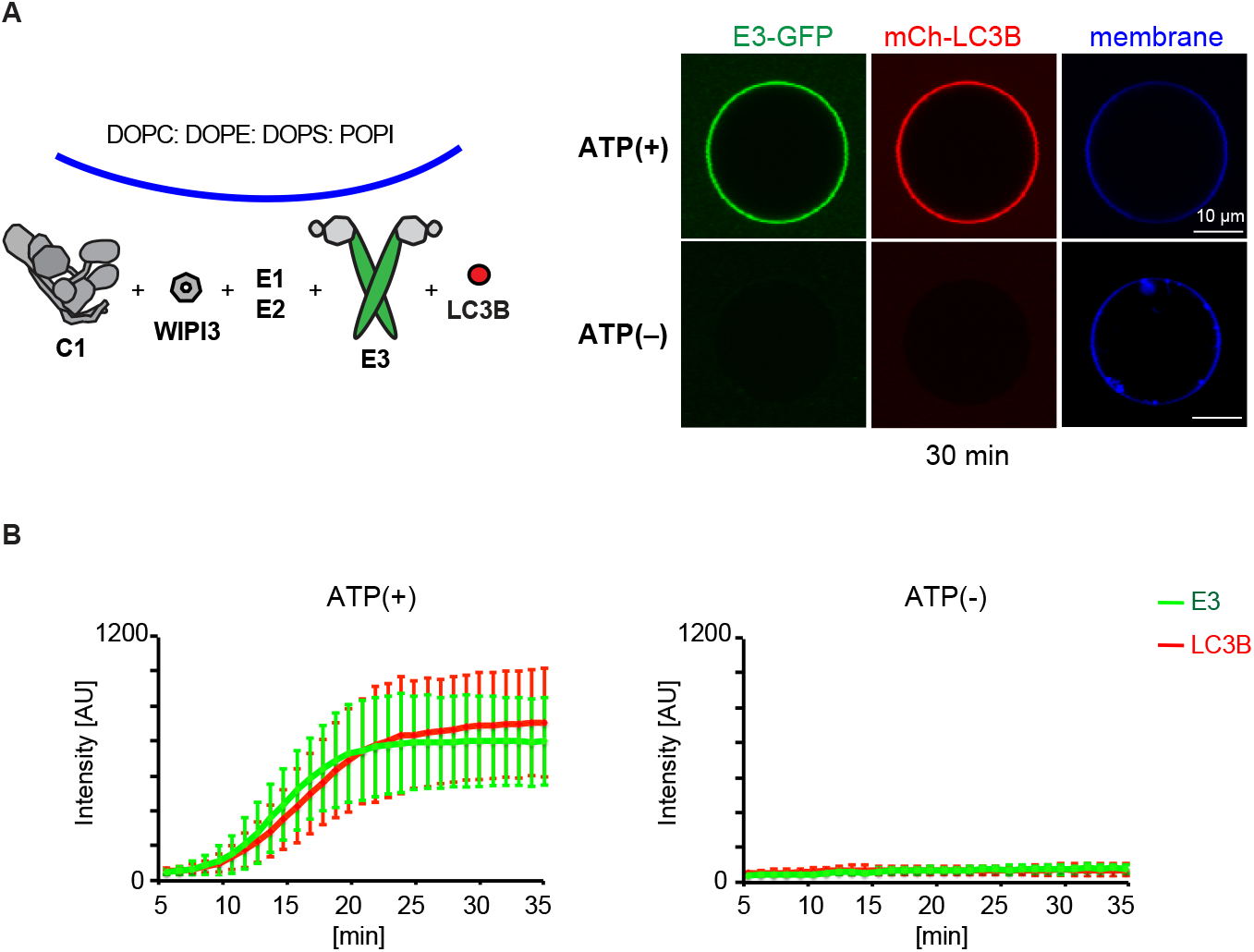
WIPI3 mediates PI3KC3-C1 trigged LC3 lipidation on GUV membranes. **A** Representative confocal images of GUVs showing E3 membrane recruitment and LC3B lipidation. mCherry-tagged LC3B was incubated with PI3KC3-C1 (100 nM), WIPI3 (400 nM), E3-GFP, ATG7, and ATG3 in the presence or absence of ATP. Scale bars, 10 μm. **B** Quantitation of the kinetics of E3 recruitment and LC3B lipidation on the membrane from individual GUV tracing in **A** (means ± SDs; N =20; 15).

## References

Anding, A.L., and E.H. Baehrecke. 2017. Cleaning House: Selective Autophagy of Organelles. Developmental Cell. 41:10–22.

Axe, E.L., S.A. Walker, M. Manifava, P. Chandra, H.L. Roderick, A. Habermann, G. Griffiths, and N.T. Ktistakis. 2008. Autophagosome formation from compartments enriched in phosphatidylinositol 3-phosphate and dynamically connected to the endoplasmic reticulum. Journal of Cell Biology. 182:685–701.

Baskaran, S., L.-A. Carlson, G. Stjepanovic, L.N. Young, D.J. Kim, P. Grob, R.E. Stanley, E. Nogales, and J.H. Hurley. 2014. Architecture and dynamics of the autophagic phosphatidylinositol 3-kinase complex. eLife. 3:e05115.

Baskaran, S., M.J. Ragusa, E. Boura, and J.H. Hurley. 2012. Two-Site Recognition of Phosphatidylinositol 3-Phosphate by PROPPINs in Autophagy. Molecular Cell. 47:339–348.

Bento, C.F., M. Renna, G. Ghislat, C. Puri, A. Ashkenazi, M. Vicinanza, F.M. Menzies, and D.C. Rubinsztein. 2016. Mammalian Autophagy: How Does It Work? Annu. Rev. Biochem.. 85:685–713.

Bigay, J., and B. Antonny. 2012. Curvature, Lipid Packing, and Electrostatics of Membrane Organelles: Defining Cellular Territories in Determining Specificity. Developmental Cell. 23:886–895.

Brier, L.W., L. Ge, G. Stjepanovic, A.M. Thelen, J.H. Hurley, and R. Schekman. 2019. Regulation of LC3 lipidation by the autophagy-specific class III phosphatidylinositol-3 kinase complex. Mol Biol Cell. 30:1098–1107.

Brier, L.W., M. Zhang, and L. Ge. 2016. Mechanistically dissecting autophagy: insights from in vitro reconstitution. Journal of Molecular Biology.

Chang, C., L.N. Young, K.L. Morris, S. von Bülow, J. Schöneberg, H. Yamamoto-Imoto, Y. Oe, K. Yamamoto, S. Nakamura, G. Stjepanovic, G. Hummer, T. Yoshimori, and J.H. Hurley. 2019. Bidirectional Control of Autophagy by BECN1 BARA Domain Dynamics. Molecular Cell. 73:339–353.e336.

Dooley, H.C., M. Razi, H.E.J. Polson, S.E. Girardin, M.I. Wilson, and S.A. Tooze. 2014. WIPI2 Links LC3 Conjugation with PI3P, Autophagosome Formation, and Pathogen Clearance by Recruiting Atg12-5-16L1. Molecular Cell. 55:238–252.

Dudley, L.J., A.G. Cabodevilla, A.N. Makar, M. Sztacho, T. Michelberger, J.A. Marsh, D.R. Houston, S. Martens, X. Jiang, and N. Gammoh. 2019. Intrinsic lipid binding activity of ATG16L1 supports efficient membrane anchoring and autophagy. The EMBO Journal. 38:e100554.

Fan, W., A. Nassiri, and Q. Zhong. 2011. Autophagosome targeting and membrane curvature sensing by Barkor/Atg14(L). Proceedings of the National Academy of Sciences of the United States of America. 108:7769–7774.

Fracchiolla, D., J. Sawa-Makarska, B. Zens, A. de Ruiter, G. Zaffagnini, A. Brezovich, J. Romanov, K. Runggatscher, C. Kraft, B. Zagrovic, and S. Martens. 2016. Mechanism of cargo-directed Atg8 conjugation during selective autophagy. eLife. 5:e18544.

Gomes, L.C., and I. Dikic. 2014. Autophagy in Antimicrobial Immunity. Molecular Cell. 54:224–233.

Gui, X., H. Yang, T. Li, X. Tan, P. Shi, M. Li, F. Du, and Z.J. Chen. 2019. Autophagy induction via STING trafficking is a primordial function of the cGAS pathway. Nature. 567:262–266.

Hanada, T., N.N. Noda, Y. Satomi, Y. Ichimura, Y. Fujioka, T. Takao, F. Inagaki, and Y. Ohsumi. 2007. The Atg12-Atg5 Conjugate Has a Novel E3-like Activity for Protein Lipidation in Autophagy. Journal of Biological Chemistry. 282:37298–37302.

Hurley, J.H., and L.N. Young. 2017. Mechanisms of Autophagy Initiation. Annu. Rev. Biochem.. 86:225–244.

Ichimura, Y., T. Kirisako, T. Takao, Y. Satomi, Y. Shimonishi, N. Ishihara, N. Mizushima, I. Tanida, E. Kominami, M. Ohsumi, T. Noda, and Y. Ohsumi. 2000. A ubiquitin-like system mediates protein lipidation. Nature. 408:488.

Itakura, E., C. Kishi, K. Inoue, and N. Mizushima. 2008. Beclin 1 Forms Two Distinct Phosphatidylinositol 3-Kinase Complexes with Mammalian Atg14 and UVRAG. Molecular Biology of the Cell. 19:5360–5372.

Itakura, E., and N. Mizushima. 2009. Atg14 and UVRAG: mutually exclusive subunits of mammalian Beclin 1-PI3K complexes. Autophagy. 5:534–536.

Itakura, E., and N. Mizushima. 2010. Characterization of autophagosome formation site by a hierarchical analysis of mammalian Atg proteins. Autophagy. 6:764–776.

Juris, L., M. Montino, P. Rube, P. Schlotterhose, M. Thumm, and R. Krick. 2015. PI3P binding by Atg21 organises Atg8 lipidation. The EMBO Journal. 34:955.

Kageyama, S., H. Omori, T. Saitoh, T. Sone, J.-L. Guan, S. Akira, F. Imamoto, T. Noda, and T. Yoshimori. 2011. The LC3 recruitment mechanism is separate from Atg9L1-dependent membrane formation in the autophagic response against Salmonella. Molecular Biology of the Cell. 22:2290–2300.

Karanasios, E., E. Stapleton, M. Manifava, T. Kaizuka, N. Mizushima, S.A. Walker, and N.T. Ktistakis. 2013. Dynamic association of the ULK1 complex with omegasomes during autophagy induction. Journal of Cell Science. 126:5224.

Kirkin, V., and V.V. Rogov. 2019. A Diversity of Selective Autophagy Receptors Determines the Specificity of the Autophagy Pathway. Molecular Cell. 76:268–285.

Koyama-Honda, I., E. Itakura, T.K. Fujiwara, and N. Mizushima. 2013. Temporal analysis of recruitment of mammalian ATG proteins to the autophagosome formation site. Autophagy. 9:1491–1499.

Krick, R., R.A. Busse, A. Scacioc, M. Stephan, A. Janshoff, M. Thumm, and K. Kuehnel. 2012. Structural and functional characterization of the two phosphoinositide binding sites of PROPPINs, a beta-propeller protein family. Proceedings of the National Academy of Sciences of the United States of America. 109:E2042–E2049.

Kuma, A., N. Mizushima, N. Ishihara, and Y. Ohsumi. 2002. Formation of the approximately 350-kDa Apg12-Apg5.Apgl6 multimeric complex, mediated by Apg16 oligomerization, is essential for autophagy in yeast. The Journal of biological chemistry. 277:18619–18625.

Lamb, C.A., T. Yoshimori, and S.A. Tooze. 2013. The autophagosome: origins unknown, biogenesis complex. Nature Reviews Molecular Cell Biology. 14:759.

Lystad, A.H., S.R. Carlsson, L.R. de la Ballina, K.J. Kauffman, S. Nag, T. Yoshimori, T.J. Melia, and A. Simonsen. 2019. Distinct functions of ATG16L1 isoforms in membrane binding and LC3B lipidation in autophagy-related processes. Nature Cell Biology. 21:372–383.

Matsunaga, K., E. Morita, T. Saitoh, S. Akira, N.T. Ktistakis, T. Izumi, T. Noda, and T. Yoshimori. 2010. Autophagy requires endoplasmic reticulum targeting of the PI3-kinase complex via Atg14L. The Journal of Cell Biology. 190:511–521.

Matsunaga, K., T. Saitoh, K. Tabata, H. Omori, T. Satoh, N. Kurotori, I. Maejima, K. Shirahama-Noda, T. Ichimura, T. Isobe, S. Akira, T. Noda, and T. Yoshimori. 2009. Two Beclin 1-binding proteins, Atg14L and Rubicon, reciprocally regulate autophagy at different stages. Nat.Cell Biol. 11:385–396.

Mercer, T.J., A. Gubas, and S.A. Tooze. 2018. A molecular perspective of mammalian autophagosome biogenesis. Journal of Biological Chemistry. jbc.R117.810366.

Metlagel, Z., C. Otomo, G. Takaesu, and T. Otomo. 2013. Structural basis of ATG3 recognition by the autophagic ubiquitin-like protein ATG12. Proc Natl Acad Sci U S A. 110:18844–18849.

Mizushima, N., A. Kuma, Y. Kobayashi, A. Yamamoto, M. Matsubae, T. Takao, T. Natsume, Y. Ohsumi, and T. Yoshimori. 2003. Mouse Apg16L, a novel WD-repeat protein, targets to the autophagic isolation membrane with the Apg12-Apg5 conjugate. Journal of Cell Science. 116:1679.

Mizushima, N., T. Noda, and Y. Ohsumi. 1999. Apg16p is required for the function of the Apg12p-Apg5p conjugate in the yeast autophagy pathway. EMBO J. 18:3888–3896.

Mizushima, N., T. Noda, T. Yoshimori, Y. Tanaka, T. Ishii, M.D. George, D.J. Klionsky, M. Ohsumi, and Y. Ohsumi. 1998. A protein conjugation system essential for autophagy. Nature. 395:395–398.

Mizushima, N., T. Yoshimori, and Y. Ohsumi. 2011. The role of Atg proteins in autophagosome formation. Annu Rev Cell Dev Biol. 27:107–132.

Obara, K., T. Sekito, and Y. Ohsumi. 2006. Assortment of phosphatidylinositol 3-kinase complexes-Atg14p directs association of complex I to the pre-autophagosomal structure in Saccharomyces cerevisiae. Molecular Biology of the Cell. 17:1527–1539.

Otomo, C., Z. Metlagel, G. Takaesu, and T. Otomo. 2013. Structure of the human ATG12~ATG5 conjugate required for LC3 lipidation in autophagy. Nature structural & molecular biology. 20:59–66.

Proikas-Cezanne, T., Z. Takacs, P. Donnes, and O. Kohlbacher. 2015. WIPI proteins: essential PtdIns3P effectors at the nascent autophagosome. J Cell Sci. 128:207–217.

Romanov, J., M. Walczak, I. Ibiricu, S. Schuchner, E. Ogris, C. Kraft, and S. Martens. 2012. Mechanism and functions of membrane binding by the Atg5-Atgl2/Atgl6 complex during autophagosome formation. EMBO J. 31:4304–4317.

Rostislavleva, K., N. Soler, Y. Ohashi, L. Zhang, E. Pardon, J.E. Burke, G.R. Masson, C. Johnson, J. Steyaert, N.T. Ktistakis, and R.L. Williams. 2015. Structure and flexibility of the endosomal Vps34 complex reveals the basis of its function on membranes. Science. 350:aac7365.

Schütter, M., P. Giavalisco, S. Brodesser, and M. Graef. 2020. Local Fatty Acid Channeling into Phospholipid Synthesis Drives Phagophore Expansion during Autophagy. Cell. 180:135–149.e114.

Stjepanovic, G., S. Baskaran, M.G. Lin, and J.H. Hurley. 2017. Vps34 Kinase Domain Dynamics Regulate the Autophagic PI 3-Kinase Complex. Molecular Cell. 67:528–534.e523.

Vanni, S., H. Hirose, H. Barelli, B. Antonny, and R. Gautier. 2014. A sub-nanometre view of how membrane curvature and composition modulate lipid packing and protein recruitment. Nature Communications. 5:4916.

Watanabe, Y., T. Kobayashi, H. Yamamoto, H. Hoshida, R. Akada, F. Inagaki, Y. Ohsumi, and N.N. Noda. 2012. Structure-based Analyses Reveal Distinct Binding Sites for Atg2 and Phosphoinositides in Atg18. Journal of Biological Chemistry. 287:31681–31690.

Weinberger, A., F.-C. Tsai, Gijsje H. Koenderink, Thais F. Schmidt, R. Itri, W. Meier, T. Schmatko, A. Schröder, and C. Marques. 2013. Gel-Assisted Formation of Giant Unilamellar Vesicles. Biophysical Journal. 105:154–164.

Weissmann, F., G. Petzold, R. VanderLinden, i. Huis, P.J. t Veld, N.G. Brown, F. Lampert, S. Westermann, H. Stark, B.A. Schulman, and J.-M. Peters. 2016. biGBac enables rapid gene assembly for the expression of large multisubunit protein complexes. Proceedings of the National Academy of Sciences. 113:E2564.

Wen, X., and D.J. Klionsky. 2016. An overview of macroautophagy in yeast. Journal of Molecular Biology. 428:1681–1699.

Wilson, Michael I., Hannah C. Dooley, and Sharon A. Tooze. 2014. WIPI2b and Atg16L1: setting the stage for autophagosome formation. Biochemical Society Transactions. 42:1327–1334.

Zachari, M., S.R. Gudmundsson, Z. Li, M. Manifava, R. Shah, M. Smith, J. Stronge, E. Karanasios, C. Piunti, C. Kishi-Itakura, H. Vihinen, E. Jokitalo, J.-L. Guan, F. Buss, A.M. Smith, S.A. Walker, E.-L. Eskelinen, and N.T. Ktistakis. 2019. Selective Autophagy of Mitochondria on a Ubiquitin-Endoplasmic-Reticulum Platform. Developmental Cell. 50:627–643.e625.

Zaffagnini, G., and S. Martens. 2016. Mechanisms of Selective Autophagy. Journal of Molecular Biology. 428:1714–1724.

Zaffagnini, G., A. Savova, A. Danieli, J. Romanov, S. Tremel, M. Ebner, T. Peterbauer, M. Sztacho, R. Trapannone, A.K. Tarafder, C. Sachse, and S. Martens. 2018. p62 filaments capture and present ubiquitinated cargos for autophagy. The EMBO Journal. 37.

Zhong, Y., Q.J. Wang, X. Li, Y. Yan, J.M. Backer, B.T. Chait, N. Heintz, and Z. Yue. 2009. Distinct regulation of autophagic activity by Atg14L and Rubicon associated with Beclin 1-phosphatidylinositol-3-kinase complex. Nature Cell Biology. 11:468–476.

